# TCRen: predicting TCR recognition of unseen epitopes based on residue-level pairwise statistical potential

**DOI:** 10.1101/2022.02.15.480516

**Authors:** Vadim K. Karnaukhov, Dmitrii S. Shcherbinin, Anton O. Chugunov, Dmitriy M. Chudakov, Roman G. Efremov, Ivan V. Zvyagin, Mikhail Shugay

## Abstract

Prediction of TCR-peptide interactions has great importance for therapy of cancer, infectious and autoimmune diseases, but remains a major challenge, particularly for unseen epitopes. We present a structure-based method that enables scoring of TCR-peptide interactions using an energy potential (TCRen) derived from statistics of TCR-peptide contacts in existing crystal structures. We show that TCRen has high performance in discriminating cognate/unrelated peptides and can facilitate the identification of cancer neoepitopes recognized by tumor-infiltrating lymphocytes.

## Main

T-cell receptor (TCR) recognition of foreign peptides (epitopes) presented by major histocompatibility complex (MHC) proteins is a crucial step in triggering the adaptive immune response. Prediction of TCR-peptide interaction is important for many clinical applications, e.g. identification of cancer neoepitopes recognized by tumor-infiltrating lymphocytes (TILs), prediction of TCR cross-reactivity for T-cell based therapy, identification of targets for antigen-specific therapy in autoimmunity, and vaccine design. Importantly, most of these tasks require predictions for novel (unseen) epitopes for which no specific TCRs are known. Several methods were developed for prediction of TCR-peptide recognition^1–4^ that utilize machine learning models of different architectures trained on data from TCR specificity databases (such as VDJdb^5^ and McPAS^6^). However, all these methods either do not support or have very low performance for unseen epitopes. To overcome this issue we developed a novel computational method, which estimates energy score of peptide-TCR interaction based on contact information extracted from TCR-peptide-MHC structures, that makes it is suitable for prediction of TCR recognition of unseen epitopes. In this manuscript we focused on the task of ranking candidate epitopes for a given target TCR, however, in principle, our method can also be applied for ranking candidate TCRs recognizing the given epitope.

The key element of our method is a statistical potential TCRen (from “TCR energy”) that describes interaction energy scores for all 380 (20×19, since cysteines do not occur in peptide-contacting regions of TCRs) possible contact pairs of TCR and peptide residues (**Figure 1a**). The procedure of TCRen derivation included the following steps: 1) all available TCR-peptide-MHC structures from Protein Data Bank (PDB)^7^ were collected; 2) a non-redundant set of these structures was selected with pairwise Levenstein distances in CDR3 regions and peptides no less 6; 3) all pairwise TCR-peptide residue contacts were extracted from the non-redundant dataset and for each of 380 possible residue pairs we calculated the “observed” number of its occurrences in the database (#*contact*(*a*, *b*), where *a* and *b* correspond to residues from TCR and peptide, respectively) and its frequency (*p_obs_* (*a*, *b*)); 4) for each of 380 residue pairs we also calculated its “expected” frequency from random pairing preferences *p_exp_* (*a*, *b*) (see **Methods** for details). Then we assumed that the more frequently the given residue pair occurs compared to its expected frequency, the more favorable interaction energy score it has, and calculated TCRen potential values for energy score of interaction between TCR and peptide residues as 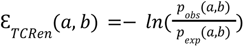.

**Figure 1.**
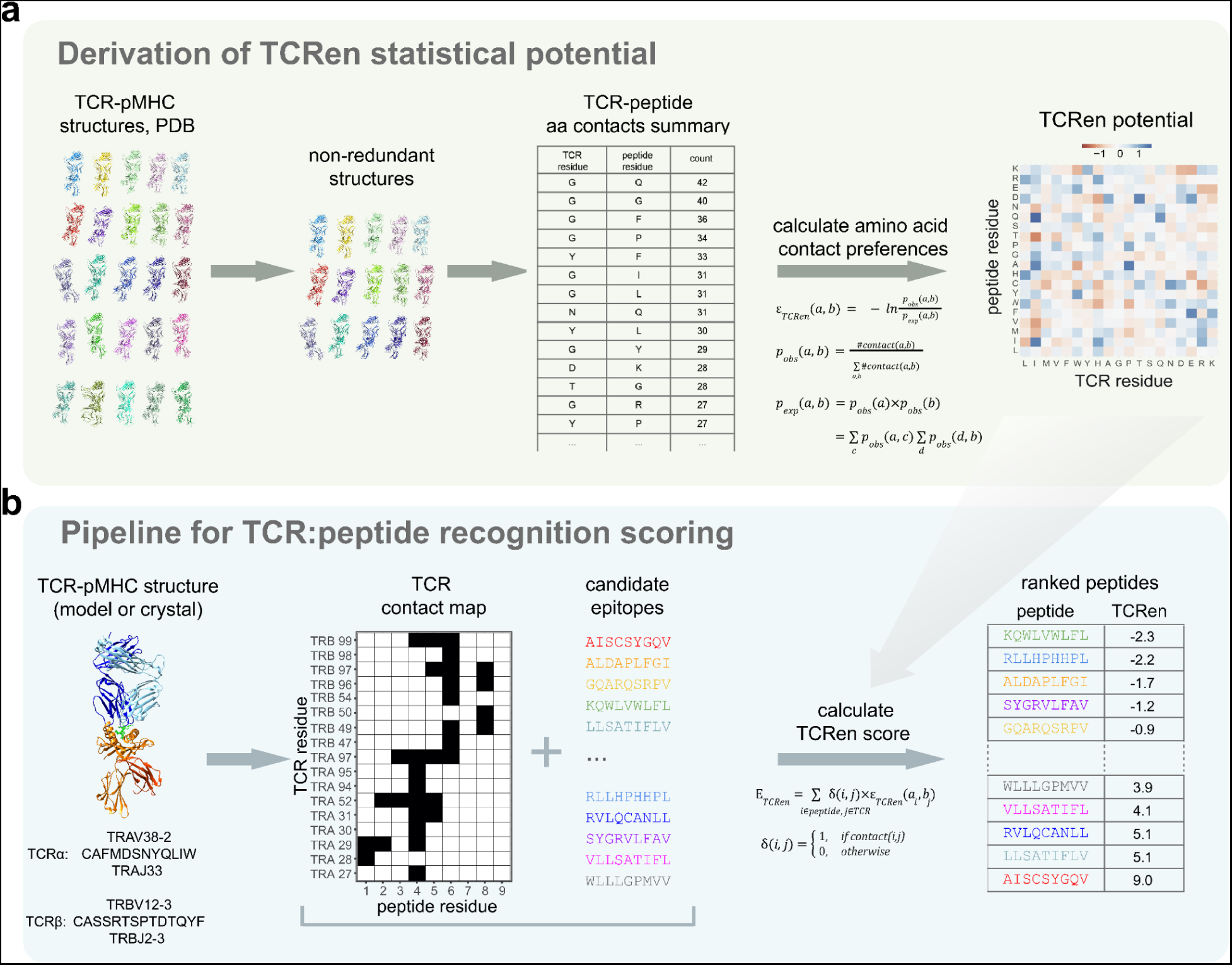
TCRen method for prediction of TCR-peptide recognition. **a.** Derivation of TCRen potential based on statistical analysis of contact preferences in TCR-peptide interactions. A non-redundant set (in terms of CDR3α, CDR3β, and peptide sequences) of TCR-peptide-MHC complexes (n = 140, which included structures with MHC class I and II from mouse and human) is selected from all available TCR-peptide-MHC crystal structures (n = 239). Contacts between TCR and peptide residues are identified and summarized (total number of contacts = 3,106). A 20 × 19 matrix of contact odds is then derived by normalizing contact frequencies to their expected frequencies from random pairing. Rows and columns in this matrix correspond to amino acids from the peptide and TCR, respectively. Cells are colored according to TCRen values. **b.** Bioinformatic pipeline for ranking of candidate epitopes, starting with a structure of the TCR-peptide-MHC complex—either experimentally derived or a homology model—followed by TCR-peptide contact map extraction and convolution with TCRen potential, leading to the final TCRen score. Our method outputs a ranked list of candidate peptides according to their estimated TCR binding energy score.

Our method starts from a structure of the peptide-MHC complex with the TCR of interest, then extracts a TCR-peptide contact map, estimates the TCRen score for all candidate peptides by convolving the contact map with TCRen pairwise energies, and finally, generates a ranked list of peptides based on this score (**Figure 1b**). The input TCR-peptide-MHC structure may be either taken from PDB (if it is available for the target TCR; e.g. for the task of prediction of cross-reactivity of a known TCR) or generated using homology modeling (in that case only sequences of TCR and candidate epitopes are required as input). The stage of homology modeling takes only several minutes (e.g. using TCRpMHCmodels^8^ software) and other stages (extraction of contacts and scoring of candidate peptides) are also performed within minutes in a simple laptop, that makes our method suitable for practical applications in which hundreds or thousands of TCR-peptide pairs are being screened.

To assess performance of our method, we first constructed a benchmark set of TCR-epitope pairs for which crystal structure of corresponding TCR-peptide-MHC complex is available. Such design of a benchmark allows to discriminate errors of TCRen potential from errors of modeling. To focus on unseen epitopes, we included in this benchmark only epitopes for which no other specific TCRs are known except the target one. As a metric of performance we used “cognate epitope rank” which we defined as a fraction of unrelated peptides (random mismatched peptides or experimentally defined binders for the same HLA allele; see **Methods** for details) which have better TCRen score than the cognate epitope. Validation was performed in a leave-one-out (LOO) setting, when for generation of predictions for each single target TCR we used TCRen version derived based on all other structures of the non-redundant set (thus, the Levenshtein distance between the derivation and test structures was no less than 6, when computed by CDR3α, CDR3β and peptide sequences). Performance of TCRen was compared with sequence-based state-of-art methods for prediction of TCR recognition of unseen epitopes: TITAN^1^ and ERGO-II^2^ (other TCR specificity prediction methods, such as NetTCR-2.0^3^ and TCRex^4^, use models separately trained for each epitope and thus do not support unseen epitopes). For comparison we also considered other structure-based energy potentials, developed for prediction of general protein-protein interactions (Keskin^9^) or protein folding (MJ^10^), and a random potential, obtained by shuffling of TCRen values. The best performance in the benchmark was demonstrated by TCRen, which had the median cognate epitope rank 19% (**Figure 2a**) and ROC AUC 0.7 (**Figure 2b**). Sequence-based methods TITAN and ERGO-II and structure-based statistical potentials MJ and Keskin performed significantly worse, having median cognate epitope ranks 34%, 40%, 40% and 54%, respectively (**Figure 2a**). We also ran the same benchmark using the full TCRen version derived based on all the structures from our non-redundant TCR-peptide-MHC dataset. The performance in this setting was better than in LOO validation, with median cognate epitope rank 4% (**Figure 2a**). This increase in performance might be explained by refinement of values of interaction energies for residues which have low frequency in CDR3 regions or epitopes. Also these results highlight the perspective to continuous increase of TCRen accuracy with the growth of the number of available TCR-peptide-MHC crystal structures.

**Figure 2.**
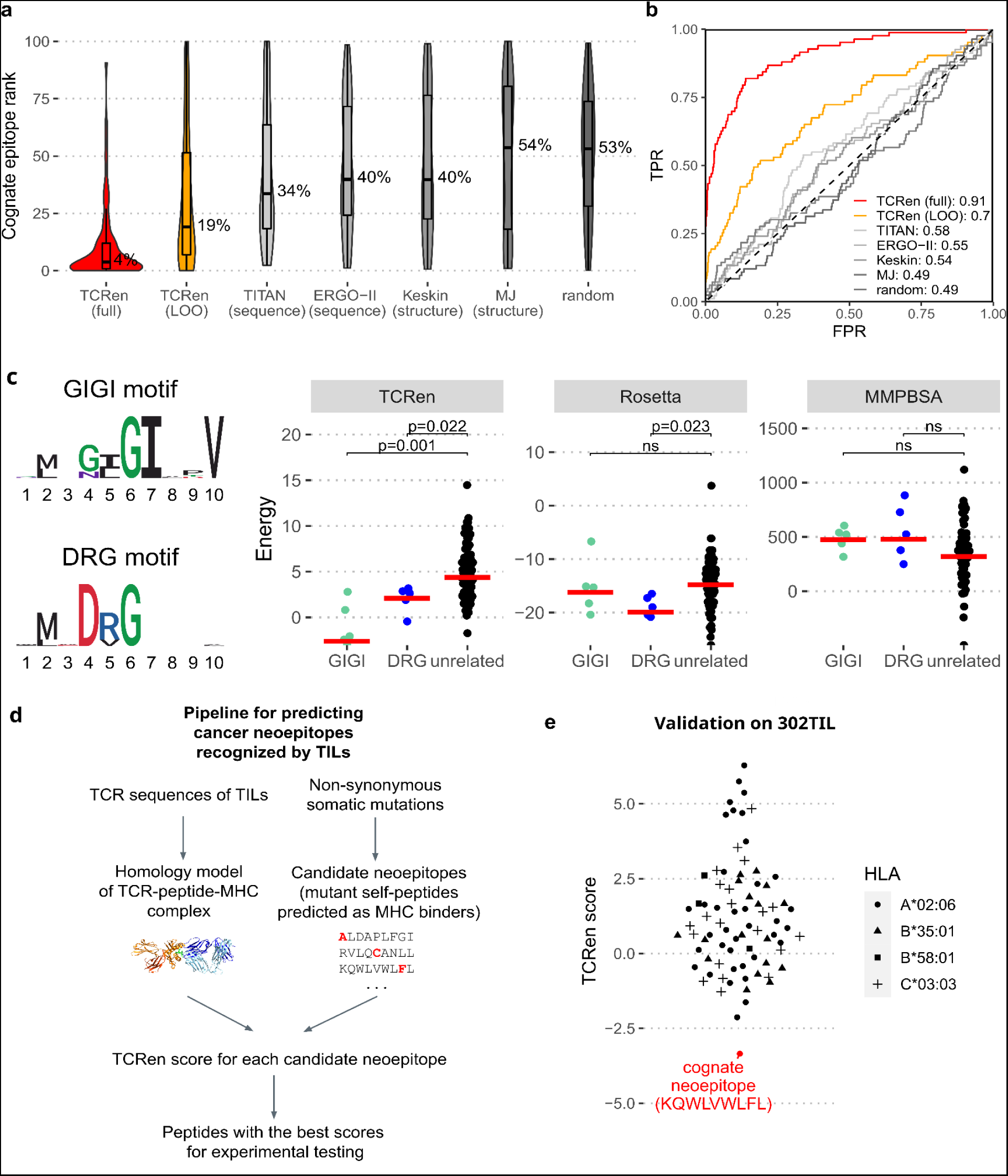
TCRen performance in distinguishing cognate TCR epitopes from unrelated peptides. **a.** Distributions of cognate epitope ranks (percentile ranks of peptide-TCR interaction energies for cognate epitopes among a set of unrelated peptides) for full version of TCRen (red), TCRen in leave-one-out (LOO) setting (orange) and other methods (shades of grey): sequence-based methods for prediction of TCR-peptide recognition TITAN^1^ and ERGO-II^2^, structure-based statistical potentials from the works of Miyazawa and Jernigan (MJ)^10^ and Keskin et al.^9^, and random potential obtained by shuffling of TCRen values. See Main text and Methods for details of the benchmarking experiment. **b.** ROC curves for distinguishing all complexes with cognate epitopes versus all complexes with unrelated peptides using the potentials mentioned in panel a. TPR = true positive rate, FPR = false positive rate. **c.** Performance of TCRen, Rosetta, and the MMPBSA approach for identifying peptides recognized by the DMF5 TCR. At left are the sequence logos for peptides with GIGI and DRG motifs recognized by DMF5 TCR. Plots show comparison of energies of interaction of DMF5 TCR with GIGI-containing (n = 5), DRG-containing (n = 5), and unrelated (n = 64) peptides as calculated using TCRen, Rosetta, and MMPBSA. Energy score calculations are averaged across 1,000 frames of 100-ns MD trajectories of the corresponding TCR-peptide-MHC complexes. **d.** Scheme of the TCRen-based bioinformatic pipeline for identifying neoepitopes recognized by tumor-infiltrating lymphocytes. **e.** TCRen values for energy score of interaction of TCR 302TIL with 79 candidate neoepitopes predicted based on exome sequencing of tumor and normal tissues in Bobisse et al.^16^. Each candidate epitope carries a somatic non-synonymous mutation and is predicted to bind to one of the patient’s HLA variants (shown as shape). Contact maps were taken from homology models of 302TIL complexes with poly-V peptide and HLA variant for which the considered neoepitope is predicted to bind. The cognate peptide recognized by TCR 302TIL is highlighted in red.

Our benchmark set included epitopes for human MHC class I, human MHC class II, murine MHC class I and murine MHC class II, and TCRen performance was consistent for all of these four groups (**Supplementary Figure 1a**). This result highlights TCRen utility for diverse HLA-restrictions and reflects the common biophysical features of TCR-peptide recognition across MHC classes and species. Also, it should be noted that 48 out of 140 (34%) structures in our non-redundant TCRen derivation set contain variants of HLA-A*02, the most characterized HLA molecule. To demonstrate that this fact does not bias TCRen predictions, we derived TCRen based exclusively on a subset of structures which contain MHC variants other than HLA-A*02 (n = 92) and then applied this TCRen version for ranking candidate epitopes for HLA-A*02-containing benchmark cases. Analogous analysis was performed for the most frequent V genes in our non-redundant dataset, TRAV12-2 (n = 16 structures) and TRBV19 (n = 12): candidate epitopes for TCRs containing these genes were ranked using TCRen version derived with exclusion of TRAV12-2 or TRBV19-containing structures. In all three experiments, for HLA-A*02 (**Supplementary Figure 1b**), TRAV12-2 (**Supplementary Figure 1c**) and TRBV19 (**Supplementary Figure 1d**), the obtained distribution of cognate epitope ranks was similar to those of TCRen in LOO setting (**Figure 2a**) that indicates that MHC and V-gene composition of the derivation set do not bias TCRen predictions.

We further tested if TCRen can be used for prediction of cross-reactivity and refinement of specificity of TCRs for which one epitope is already known. For this benchmark we used data from the work of Birnbaum *et al.*^11^, in which the authors utilized yeast display screening to identify peptides recignized by three distinct TCRs—2B4, 226, 5cc7—restricted to murine class II MHC I-E^k^. We tested the ability of TCRen to distinguish between peptide which are either recognized (top-20 yeast display hits with the highest enrichment after 5 rounds of selection) or unrelated (predicted I-E^k^ binders). For candidate epitope scoring we used the version of TCRen potential derived without the structures containing TCRs similar to 2B4, 226 and 5cc7. For all three TCRs, TCRen demonstrated good performance, with an AUROC of 0.98, 0.74, and 0.74 for 2B4, 226, and 5cc7, respectively (**Supplementary Figure 2**).

Next, we compared the performance of TCRen with fine-grained atomic-level methods that are widely used for the assessment of protein-protein interactions: Rosetta scoring function^12^ and the molecular mechanics Poisson-Boltzmann surface area (MMPBSA) approach^13^. Since atom-level calculations of Rosetta and MMPBSA require generation of ensembles of structures using highly computational expensive molecular dynamics (MD) simulations, we performed this comparison on a single example of DMF5 TCR. We selected this case as this TCR has an interesting specificity: its cognate epitopes that were previously identified by yeast display^14^, are clustered into two groups, one with a hydrophobic core (GIGI in positions 4–7), and the other with a charged core (DRG in positions 4–6) (**Figure 2c**, left panel). We performed MD simulations of DMF5 TCR-peptide-MHC complexes with experimentally-confirmed epitopes containing either GIGI or DRG motifs as well as unrelated peptides having neither of these motifs. Then we calculated the trajectory-averaged energy score of TCR-peptide interaction using TCRen, Rosetta, and MMPBSA. TCRen was the only approach to report lower (more favorable) energy score values for both GIGI- and DRG-containing peptides compared to unrelated peptides. The Rosetta scoring function assigned lower energy score values for DRG but not for GIGI peptides, while the MMPBSA approach showed no significant differences between unrelated and cognate peptides from both groups (**Figure 2c**, right panel). The MD analysis also confirmed the robustness of our TCRen calculations which comes from its residue-level definition in contrast to atom-level Rosetta and MMPBSA (**Supplementary Figure 3**).

According to our benchmarks, TCRen has a good performance in prediction of TCR-peptide recognition, while methods developed for general protein-protein interactions fail in this task. To explain this, we compared matrices of TCRen, MJ and Keskin potentials (**Supplementary Figure 4a**). MJ and Keskin have similar patterns with the clustering of non-polar, polar and charged residues (**Supplementary Figure 4a**) with the best scores for contacts of non-polar residues and the worst scores for charged residues (**Supplementary Figure 4b**). TCRen doesn’t have a simple regular pattern analogous to that of MJ and Keskin. Instead, TCRen reflects the unique nature of TCR-peptide recognition with a higher impact of medium-strength, but specific contacts mediated by polar amino acids (rather than non-specific hydrophobic interactions which are predominant in protein-protein and intraprotein interactions). In the same time, TCRen captures basic biophysical principles: TCRen score is favorable for interactions between the residues with the opposite charges and disfavorable if the charges are the same (**Supplementary Figure 4c**). We also performed analysis of TCRen, MJ and Keskin potentials’ correlations with physicochemical properties of contacting amino acids. The results indicated that TCRen accounts for complex interactions of physicochemical properties of contacting amino acids in contrast to MJ and Keskin potentials mostly governed by hydrophobic interactions (see **Supplementary Note 1**). Also, TCRen reflects the asymmetry of TCR-peptide interfaces with depletion of hydrophobic residues and enrichment of polar amino acids in CDR3 compared to MHC-presented peptides (**Supplementary Figure 5**) which can be attributed to the thymic depletion of TCRs with highly hydrophobic CDR3s that are highly cross-reactive due to their ability to form non-specific hydrophobic interactions^15^.

In practical applications, crystal structure for the target TCR is not available, so homology modeling of TCR-peptide-MHC complex should be used. That raises the question if the accuracy of existing modeling methods can enable reliable TCRen predictions. To test this, we assessed TCRen performance when homology models are used as input. For homology modeling we used TCRpMHCmodels software. We controlled that the target structures were not used as templates, validation was performed using LOO procedure. The performance in the setting with homology models was only slightly lower compared to the results obtained with crystal structures used as input (**Supplementary Figure 6a**). The performance was still good when the best available templates for modeling had sequence identity only 50-65% to the target one which may be regarded as a typical value in practical applications (**Supplementary Figure 6b**). We interpret these results as an indication that the energy score of interaction between TCR and its cognate epitope is robust to deviation from the exact orientation (seen in crystal structure). From a biological state of view, a non-perfect homology model may be regarded as mimicking the first stages of *in vivo* antigen recognition when the docking mode of TCR has not still been adapted to the optimal one but TCR affinity should already be high enough to maintain the interaction.

TCRen can be used for a variety of applications in investigation and treatment of cancer, autoimmunity and infectious diseases, e.g. for the identification of cancer neoepitopes recognized by tumor-infiltrating lymphocytes (TILs) using the bioinformatic pipeline shown in **Figure 2d**. To demonstrate it, we first focused on the data from the study of Bobisse *et al.*^16^ on TILs with TCR 302TIL from a patient with recurrent advanced epithelial ovarian cancer. Using exome sequencing of normal and cancer tissues, the authors predicted 79 candidate neoepitopes for 302TIL TCR which are peptides with nonsynonymous somatic mutations predicted as binders to HLA variants of the patient. All these candidate neoepitopes were experimentally tested and one of them (KQWLVWLFL presented by HLA-A*02:06) was validated as a cognate neoepitope of 302TIL TCR. Since the experimental testing of all candidate epitopes is cost- and time-consuming, we examined if TCRen is useful in shortening of the list of peptides for validation. This benchmark was performed in a real-life setting, when no prior knowledge about 3D structure and HLA-restriction of TCR is available. First we generated homology models of 302TIL complexes with random peptides presented by all the patient’s HLA (structures with 302TIL were not used as templates; see **Supplementary Figure 7a**). Then we ranked all the candidate neoepitopes using the pipeline from **Figure 1b** with the homology models as inputs (see **Methods** for the details). TCRen ranked the cognate epitope KQWLVWLFL as Top-1 (**Figure 2e**). Such high performance was obtained even regardless of the fact that the used homology model shared only 60% of TCR-peptide contacts with the crystal structure.

To test TCRen accuracy for ranking candidate epitopes for TILs on a larger scale, we used the data from Bigot et al.^17^, where cognate neoepitopes of 14 cancer-reactive T cells were identified among 44 candidate neoepitopes derived from *SF3B1*^mut^-modified intron–exon junctions in uveal melanoma. TCRen ranked the cognate epitope in top-9 peptides with the best score in 6/14 cases and in top-16 peptides in 11/14 cases (**Supplementary Figure 8**), even though for the majority of these TCRs only templates with low sequence identity were available (and consequently, low quality of the models is expected). These results highlight that TCRen can be effectively used for the reduction up to several folds of the list of candidate neoepitopes for the experimental testing.

In summary, our method TCRen enables discrimination of cognate TCR epitopes from unrelated peptides using both experimental and modelled TCR crystal structures. Our results suggest that TCRen can find applications in cancer immunotherapy, autoimmunity studies, and vaccine design, and in particular for ranking candidate neoepitopes for TILs. TCRen demonstrated superior performance compared to structure-based approaches developed for general protein-protein interactions, that highlights the need to account for specific and unique nature of TCR-peptide interfaces. Also, TCRen significantly outperforms state-of-art sequence-based methods for TCR specificity prediction. This result indicates that the use of 3D structural information about pairwise TCR-peptide contacts can be a a promising way to make generalizable predictions for unseen epitopes. In the current work, we use homology models of TCR-peptide-MHC complexes to extract the contact maps. In the future, approaches based on AlphaFold^18, 19^ or geometric deep learning models^20, 21^ may also be adapted to predict TCR-peptide contacts which can further used for TCRen predictions. Thus, we expect synergistic interaction between TCRen and deep learning-based methods to enable accurate predictions of TCR specificity.

## Acknowledgements

Supported by the grant from the Ministry of Science and Higher Education of Russian Federation (075-15-2019-1789). MD simulations were supported by the HSE University Basic Research Program and carried out with the use of computational facilities of the Supercomputer Center “Polytechnical” at the St. Petersburg Polytechnic University.

## Methods

### Selection of a non-redundant set of TCR-peptide-MHC structures

All available crystal structures of TCR-peptide-MHC complexes (n = 239, **Supplementary Table 1**) were downloaded from PDB, and a non-redundant set of them was composed based on similarity of CDR3α, CDR3β, and peptide sequences as described below. For each pair of TCR-peptide-MHC crystal structures from PDB we calculated Levenshtein distance between their CDR3α, CDR3β, and peptide sequences (*d_CDR3α_, d_CDR3β_ and d_peptide_*, respectively, and the overall sequence distance *d* between these two structures was calculated as:

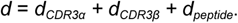

We then performed hierarchical clustering of PDB structures based on this measure, cutting the resulting tree at height = 6. The selection of this value was based on our observation from exploration of VDJdb database that pairwise Levenstein distances between CDR3 (*d_CDR3α_ + d_CDR3β_*) equal to 6 may be regarded as a threshold: pairs of sequences with lower *d* are almost never found to recognize different epitopes. Such a stringent value of cutoff ensures that all the structures in the non-redundant dataset significantly differ from each other. The results of the work are robust to the selection of the exact value of this threshold.

This procedure yielded 140 clusters. We selected a single representative from each group to include in the non-redundant set of TCR-peptide-MHC structures (n = 140, **Supplementary Table 2**). Thus, all the structures in the non-redundant set have pairwise *d* no less than 6.

### Derivation of TCRen potential and calculating TCRen score

For all the structures in the non-redundant set of TCR-peptide-MHC complexes, we extracted contacts between TCR and peptide residues. Residues were considered to be in contact if any two atoms of these residues were separated by a distance ≤ 5 Å. The number of contacts that CDR and framework regions of TCR form with peptides in different crystal structures is shown in **Supplementary Figure 9**, the number of contacts that residues in specific positions in CDRs of different lengths form with peptides is shown in **Supplementary Figure 10**.

Derivation of TCRen potential is based on the assumption that the distribution of frequencies of pairwise amino acid contacts can be described using the inverse Boltzmann relationship, which implies that the more favorable energy score of interaction ε*_TCRen_*(*a*, *b*) between residue *a* from TCR and residue *b* from peptide is, the higher frequency is has compared to expectations from random contact preferences (when ε*_TCRen_*(*a*, *b*) = *const*):

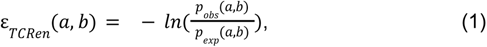

where *p_obs_*(*a*, *b*) is the observed frequency of contacts between TCR residue *a* and peptide residue *b* in the database, and *p_exp_*(*a*, *b*) is the expected frequency of contacts between these residues when assuming random pairing of amino acids.

*p_obs_*(*a*, *b*) and *p_exp_*(*a*, *b*) are calculated using the following equations:

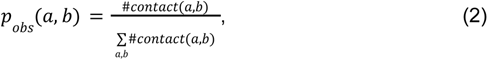

where #*contact*(*a*, *b*) is the number of contacts between residues *a* and *b* observed in the database plus pseudocount 1, and

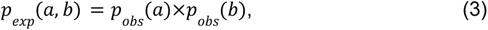

where *p_obs_*(*a*) and *p_obs_*(*b*) are the frequencies of residues *a* and *b*, respectively, in the reference state.

We compared several approaches for the selection of reference state (see below) and the best performance was achieved when using contact-fraction reference state with *p_obs_*(*a*) and *p_obs_*(*b*) calculated as:

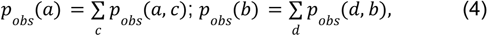

where summation is carried out through all 20 amino acids denoted with indices *c* and *d*.

The interaction energy between TCR and peptide in the given complex is estimated as:

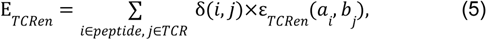

where summation is carried out through all residues *i* in the peptide and all residues *j* in the TCR; δ(*i*, *j*) = 1 if residues *i* and *j* are in contact and = 0 otherwise.

### Calculation of asymmetrical and symmetrical versions of TCRen for different reference states

We also explored other approaches for calculating TCRen. We considered three different reference states—contact-fraction, mole-fraction, and normalized mole-fraction—and derived symmetric and asymmetric versions for each of these. In the symmetric version, we considered contacts without assignment of which molecule type (peptide or TCR) the contacting residues come from:

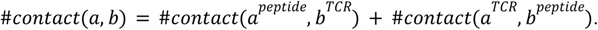

That resulted in

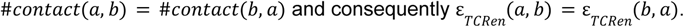

In the asymmetric version, we explicitly considered chain types corresponding to both interacting residues:

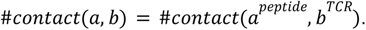

In general case, this resulted in:

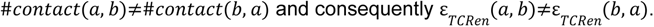

In the asymmetric version, *p_obs_*(*a*) and *p_obs_*(*b*) are also calculated separately for peptide and TCR, and equation (3) may be rewritten as:

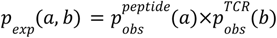

Calculation of *p_exp_*(*a*, *b*) depends on the selected reference state. We tested three possible reference states:

1. Contact-fraction, in which *p_obs_*(*a*) and *p_obs_*(*b*) are calculated as the aggregated fraction of contacts that residue *a* or *b* forms with all other residues:

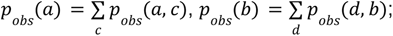
2. Mole-fraction, in which database: *p_obs_*(*a*) is the mole fraction of residue *a* in all peptides in the database:

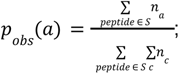

and *p_obs_*(*b*) is the mole fraction of residue *b* in all CDR3s in the database:

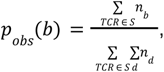

where *n_i_* is the number of occurrences of residue *i* in the given peptide/CDR3 and summation is carried out through all peptides/CDR3s in database *S*;
3. Normalized mole-fraction, in which residue frequencies are normalized on the contact for residues in the given position of TCR/peptide (**Supplementary Figure 10**):

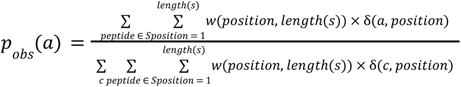 And 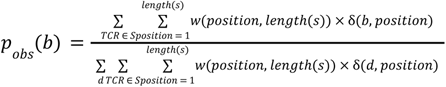, where *w*(*position*, *length*) is the average number of contacts that a residue in a given position of peptide/CDR3 of the given length forms with its counterpart. δ(*i*, *position*) = 1 if there is residue *i* in the given position of peptide/CDR3, and = 0 if there is another residue in this position.

Note, that the mole-fraction reference state is a special case of the normalized mole-fraction reference state when stating *w* = 1 for all positions of peptide/CDR3 of all lengths.

The reason for introducing a normalized mole-fraction reference state is the different propensities of forming contacts for residues at different positions in peptides and CDRs of different lengths. As can be seen from **Supplementary Figure 10**, residues that are close to the center of peptide or CDR3 tend to form more contacts. However, substantial variation in the number of contacts for given peptide and CDR3 positions exists between structures, so the observation that a residue in the center of the peptide or CDR3 does not form contacts with its counterpart in the given structure may indicate that all available contacts are energetically disfavored and that it is more favorable to avoid any contact.

In calculating mole fractions of residues in TCRs, we considered only residues in CDR3, with trimming of the first three and the last four residues, as these residues in CDR3 almost never contact the peptide in available TCR-peptide-MHC crystal structures (**Supplementary Figure 10**). Although CDR1α/β and CDR2α/β often form a number of contacts with the peptide (**Supplementary Figure 9**), position-specific interaction patterns are not regular for these regions, and the median number of contacts is 0 for almost all residue positions in CDR1α/β and CDR2α/β (**Supplementary Figure 10**).

We compared performance of different versions of TCRen in distinction between cognate epitope and unrelated peptides in the same benchmark setting as in **Figure 2a**. The best performance was demonstrated by the asymmetric version of TCRen with contact-fraction reference state (**Supplementary Figure 11**). In the main text we consider only this version of TCRen and refer to it as TCRen.

### Cognate epitope identification among unrelated peptides

To assess performance of different methods (including TCRen) in distinguishing between unseen cognate epitope and unrelated peptides, we use “cognate epitope rank”, which is defined as the fraction of unrelated peptides which have better TCR-peptide interaction score than the cognate epitope.

For the first benchmark, in order to assess performance of TCRen potential itself and remove bias coming from modeling inaccuracies, we considered only TCR-epitope pairs for which corresponding crystal structure is available. To focus on unseen epitopes, we considered only epitopes for which no other specific TCRs are documented in the VDJdb database except the target one.

TCRen scores were calculated using the pipeline shown in **Figure 1b**. As input, we used either a crystal structure (results are presented in **Figure 2a**) or a homology model with a poly-V peptide obtained using TCRpMHCmodels software^8^ (results are presented in **Supplementary Figure 6**; in that setting we considered only the cases which are not included in TCRpMHCmodels template database and manually checked the templates used for modeling to ensure that the similarity of target TCR to the most similar template was lower than 81%).

For each entry in our benchmark (a pair of a TCR and its cognate epitope), we composed a list of 1,000 unrelated peptides with the same length and anchor residues as the cognate epitope, but mismatched from it in all non-anchor positions. The second and last residues were considered as anchors, as they are mostly responsible for MHC binding. For calculating TCRen score for each unrelated peptide, we used contact maps extracted from the corresponding crystal structure with residues of the cognate epitope substituted with residues located at the same positions in the unrelated peptide.

For each peptide (cognate epitope and unrelated peptides) we calculated TCRen score using equation (5). Then for each structure in our non-redundant set we calculated cognate epitope rank - the fraction of unrelated peptides which have TCRen score lower (better) than the score of the cognate epitope.

We also considered another approach to the construction of the unrelated peptides reference set, using peptides from the Immune Epitope Database^22^ (IEDB) with the binding to the corresponding MHC variant confirmed by mass-spectrometry assays. In this setting we obtained almost the same distribution of cognate epitope ranks for TCRen (**Supplementary Figure 12**), highlighting that TCRen potential reflects features of TCR-peptide recognition, but not peptide presentation by MHC.

### Benchmark with yeast display data

For the yeast display benchmark, data from the study of Birnbaum *et al.*^11^ was used. This study describes yeast display screening of peptide epitopes for 2b4, 226 and 5cc7 TCRs, restricted to murine class II MHC I-E^k^. After each of five rounds of enrichment, the authors performed sequencing of yeast library. For our benchmark, for each of three TCRs we composed a positive set of 20 peptides which had the highest enrichment after the fifth round of yeast display screening. Negative set included 202 peptides which were predicted as binders to murine class II MHC I-E^k^ by NetMHCIIpan-4.0 software^23^ out of 250000 13-mer peptides with completely randomized sequence. NetMHCIIpan-4.0 was run with the default parameters and peptides with Rank < 5% were considered as binders. All peptides both from positive and negative set had the same length of 13 amino acids. TCRen scores were calculated for all the peptides and ROC curves for distinction between peptides from the positive and the negative sets were drawn. Contact maps for TCRen scoring were taken from crystal structures with PDB ID 3QIB, 3QIU and 4P2R for 2b4, 226 and 5cc7 TCRs, respectively. This setting corresponds to a user scenario when a single epitope is already known for the target TCR (in this case, ADLIAYLKQATKG for 2b4 and 226 TCR and ANGVAFFLTPFKA for 5cc7 TCR) and the task is to predict cross-reactivity to the other peptides.

### Molecular dynamics simulations of DMF5-peptide-HLA-A02 complexes

MD simulations were performed for DMF5-peptide-HLA-A02 complexes with different peptides from three groups: experimentally confirmed epitopes with GIGI motif (n = 5), experimentally confirmed epitopes with DRG motif (n = 5), and unrelated peptides (n = 64). For each of these, 100-ns MD simulations were carried out in at least three replicas for GIGI and DRG peptides and two replicas for unrelated peptides (total simulation time was 17.8 μs). Sequences for GIGI and DRG peptides were taken from Riley *et al.*^24^; and recognition by DMF5 in the HLA-A02 context was confirmed using surface plasmon resonance (SPR) and interferon γ (IFNγ) release assay. 32 unrelated peptides were taken from the IEDB database^22^ with the following conditions: they are experimentally determined to be HLA-A*02:01 binders, derived from human proteins, and contain neither GIGI nor DRG motifs. Starting structures for MD simulations were constructed based on PDB crystal structure templates (6AM5 for GIGI and 6AMU for DRG), with corresponding peptide sequence introduced by the “mutate_residue” PyRosetta protocol.

Preparation of the system and MD simulations were carried out using GROMACS v.5.1.4 software with the Amber99sb-ildn force field. The starting structure was placed in a rectangular box filled with explicit TIP3P water, with a size that allowed the distance between the complex and the box boundaries to be no less than 1.2 nm. Then the system was neutralized by addition of a necessary number of Na^+^ ions, minimized by a steepest descent method, and heated to 298 K with positional restraints for all heavy atoms. MD replicas were performed from the structures obtained after heating using different seeds for initial velocities of atoms. In order to prevent rotation of the complex leading to its interaction with periodic images, we used positional restraints on Cα-atoms of MHC residues 182 and 197 along the two smaller axes of the box, with force constants 10-fold smaller than the ones used at the heating stage. These residues were chosen as they are located in the MHC α3-domain, and so these restraints should have no impact on TCR-peptide-MHC interface behavior. The obtained trajectories were visually checked in the absence of interactions of the complex with its periodic images. Use of the rectangular box with imposed positional restraints resulted in a five-fold reduction of computational time compared to simulation of the same systems in a dodecahedronic box.

The Rosetta Interface analyzer module was used to calculate the binding energy score (dG_separated) according to the Rosetta scoring function^12^ between TCR and peptide. Molecular mechanics Poisson–Boltzmann surface area (MMPBSA) was applied for the estimation of binding energy between the TCR and peptide-MHC using g_mmpbsa tool^13^. 100-ns MD trajectories were split into 1,000 frames (snapshots), and for each frame we calculated Δ*G* using TCRen, Rosetta scoring function or MMPBSA. Finally, Δ*G* was averaged over all the frames.

### Comparison with other state-of-art methods for prediction of TCR-peptide recognition

We compared TCRen performance with two other state-of-art alternative methods for prediction of TCR specificity for comparison with TCRen: TITAN^1^ and ERGO-II^2^. Both of these methods are sequence-based and use neural networks of different architectures trained on curated databases of TCR sequences with known antigen specificities (VDJdb^5^ and McPAS-TCR^6^) and use explicit encoding of both TCR and peptide sequences that make them suitable for making predictions for unseen epitopes for which no specific TCRs are known. Other published methods for prediction of TCR specificity, such as NetTCR-2.0^3^, TCRex^4^ and tcrdist3^25^, were not included in the comparison, as they either use models trained for each epitope separately (NetTCR-2.0 and TCRex) or rely on similarity with other TCRs with known specificity (tcrdist3), that makes them unsuitable for unseen epitopes. For TITAN and ERGO-II we assessed performance on the same benchmark as we used previously for TCRen, calculating cognate epitope rank among peptides with random sequences. TITAN and ERGO-II were run according to the recommendations of the authors with the default parameters. Notably, for TITAN and ERGO-II we used the models pre-trained by the authors on TCR specificity data (from VDJdb/McPAS-TCR databases), so most entries from the benchmark test set (except most recently deposited) was also present in the training sets for these methods, that could lead to overestimation of their performance.

We also compared TCRen performance with structure-based statistical potentials, derived based on contact preferences in protein folding (from the work of Miyazawa and Jernigan (MJ)^10^) and in general protein-protein interfaces (from the work of Keskin *et al.*^9^). For this, we used the same benchmark and the same pipeline for extraction of contacts and estimation of TCR-peptide energy score of interaction as for TCRen (**Figure 2b**), but using MJ and Keskin instead of TCRen potential.

### Identification of cancer neoepitopes recognized by tumor-infiltrating lymphocytes

To demonstrate how TCRen can be applied to identify neoepitopes recognized by TILs we screened for previously published studies on neoepitope-specific TCRs in which the following information is available: 1) a list of candidate neoepitopes which were experimentally tested; 2) the crystal structure of TCR-peptide-MHC complex with the cognate neoepitope. The latter is important to discriminate between errors coming from inaccuracies of the TCRen potential itself and the errors from the modeling procedure. Both types of information were available for 302TIL TCR identified in the work of Bobisse *et al.*^16^ in a patient with ovarian cancer. According to exome sequencing of normal and tumor tissue, the same patient also featured 79 candidate neoepitopes having nonsynonymous somatic mutations and binding to one of HLA alleles of the patient. This list of candidate neoepitopes included 9-mers restricted to HLA-A*02:06, HLA-B*35:01, HLA-B*58:01 and HLA-C*03:03 and 10-mers restricted to HLA-A*02:06, HLA-B*35:01, HLA-C*03:03. Using TCRpMHCmodels^8^ software we generated homology models of 302TIL TCR complexes with poly-V peptides (9-mer VVVVVVVVV or 10-mer VVVVVVVVVV) and HLA-A*02:06, HLA-B*35:01, HLA-B*58:01 and HLA-C*03:03. Poly-V was chosen for modeling because valine has an intermediate value of side-chain volume amongst the 20 canonical amino acids. TCRen score was then calculated using contact maps of the homology models. Example templates produced by TCRpMHCmodels for 302TIL-VVVVVVVVV-HLA-A*02:06 are shown in **Supplementary Figure 7A**. To examine the accuracy of the obtained homology model, we superimposed it with the crystal structure of 302TIL with its cognate neoepitope presented by HLA-A*02:06, taken from the study of Devlin *et al.*^26^ (PDB ID: 6UK4; **Supplementary Figure 7B**).

To demonstrate the utility of TCRen to identify neoepitopes recognized by tumor-reactive TILs on a larger scale, we used the data from Bigot et al.^17^, where 44 HLA-A*02:01-restricted 9-mer neoepitopes derived from abnormal junctions in *SF3B1*^mut^ uveal melanoma were predicted and tested for recognition by 18 tumor-reactive T cell clones. We excluded 4 TCRs with CDR3β length more than 16 amino acids due to limitations of existing modeling software in predicting long loops. For the remaining 14 TCRs we performed homology modeling of TCR-peptide-MHC complexes with 9-mer VVVVVVVVV peptide and HLA-A*02:01 and ranked 44 candidate neoepitopes according to the TCRen score using the pipeline from **Figure 1b**.

## Code availability

All the code and data required to reproduce the analysis performed in the study, as well as script and tutorial for running TCRen on new data are available at GitHub repository https://github.com/antigenomics/tcren-ms.

## Supplementary Materials

### Supplementary Figures

**Supplementary Figure 1.**
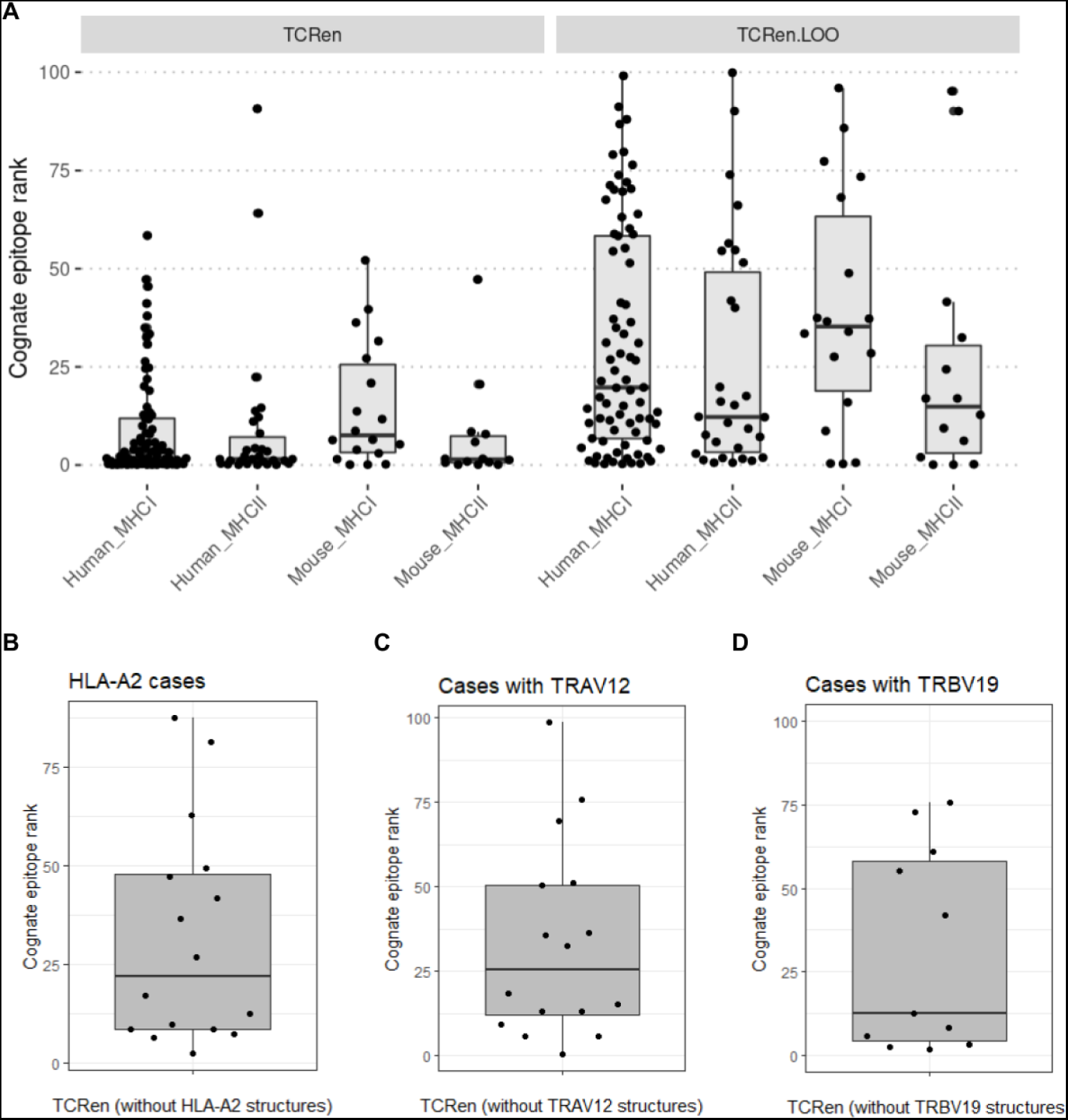
Performance of TCRen in distinction between cognate epitope and unrelated peptides for different subsets of benchmark cases. **a.** Cognate epitope ranks for complexes with MHC class I (MHCI) or class II (MHCII) from human or mouse. **B.** Cognate epitope ranks for HLA-A2-containing benchmark cases with unseen epitopes, calculated using TCRen version derived without HLA-A2-containing structures. **C-D.** Cognate epitope ranks for TRAV12-2- (**C**) and TRBV19-containing (**D**) benchmark cases, calculated using TCRen version derived without TRAV12-2- (**C**) and TRBV19-containing (**D**) structures.

**Supplementary Figure 2.**
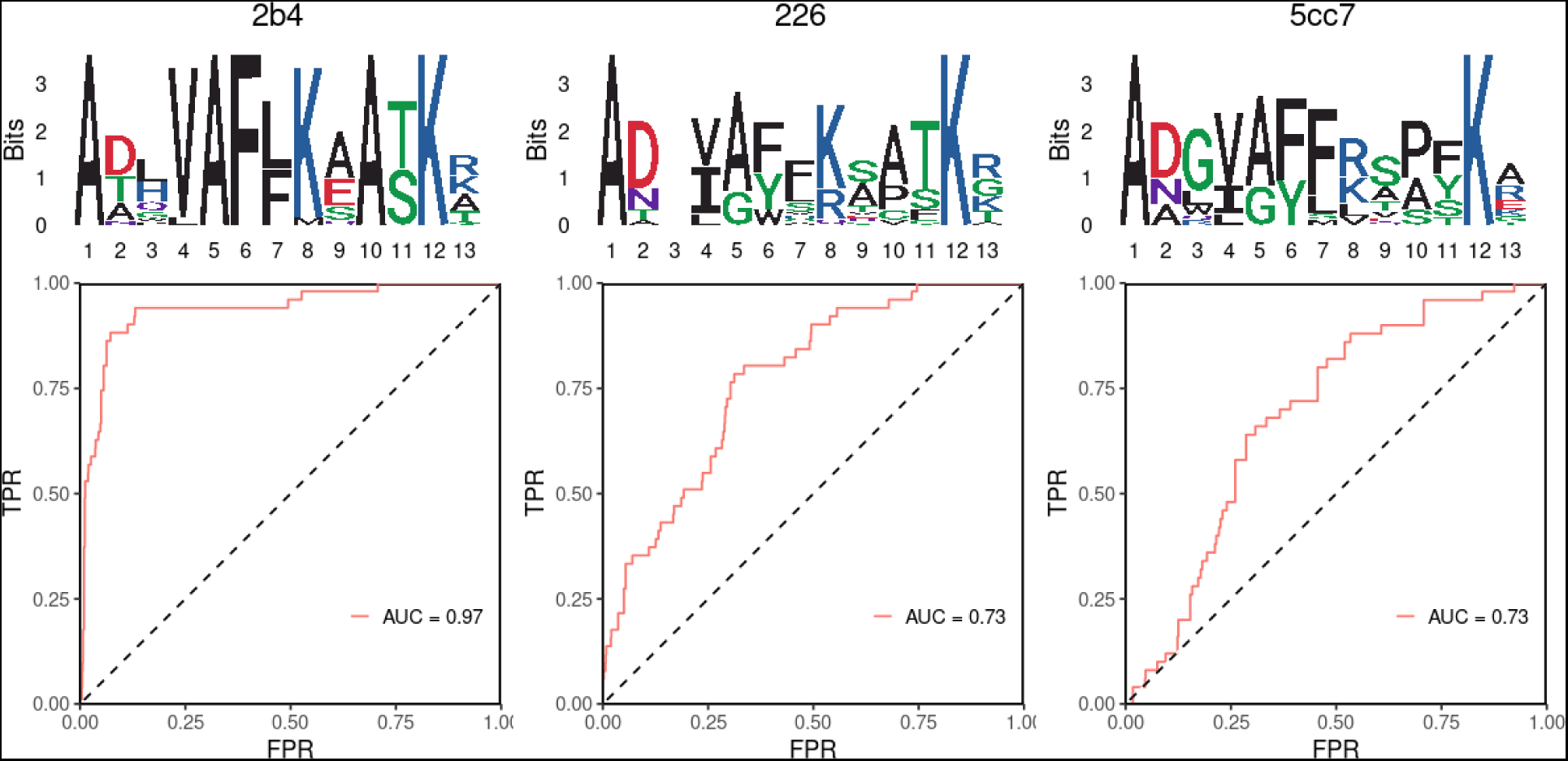
Performance of TCRen in discriminating a set of peptides recognized by TCRs 2B4, 226, and 5cc7, which were previously identified from yeast display screening^11^, and unrelated peptides. Top shows sequence logos for the yeast display hits, bottom shows ROC curves for 2B4, 226, and 5cc7.

**Supplementary Figure 3.**
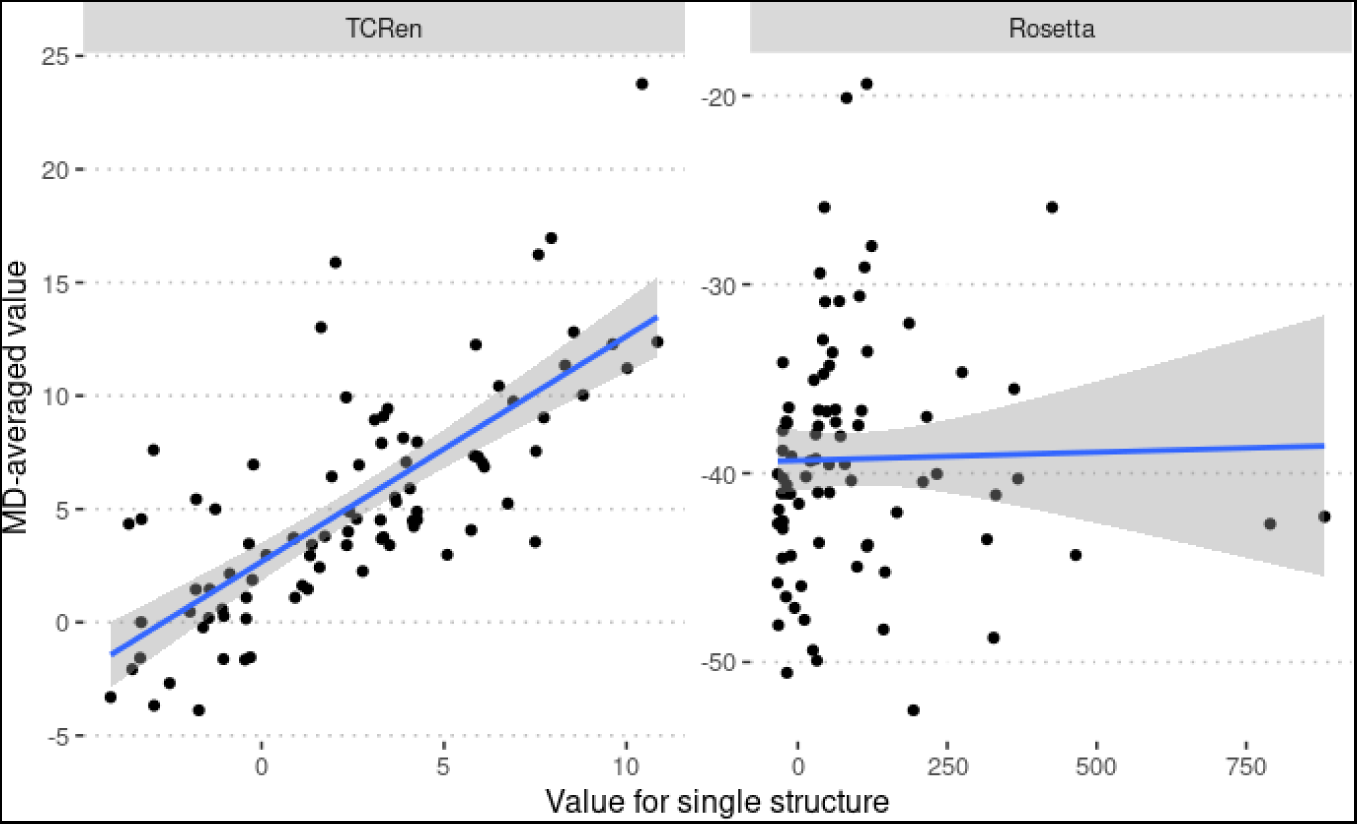
Robustness of TCRen and Rosetta potentials. Y axis correspond to values of TCRen/Rosetta scores (energy score of TCR-peptide interaction) averaged across 1000 frames of 100-ns MD trajectories, X axis - values calculated from one static structure obtained using “fixbb” protocol.

**Supplementary Figure 4.**
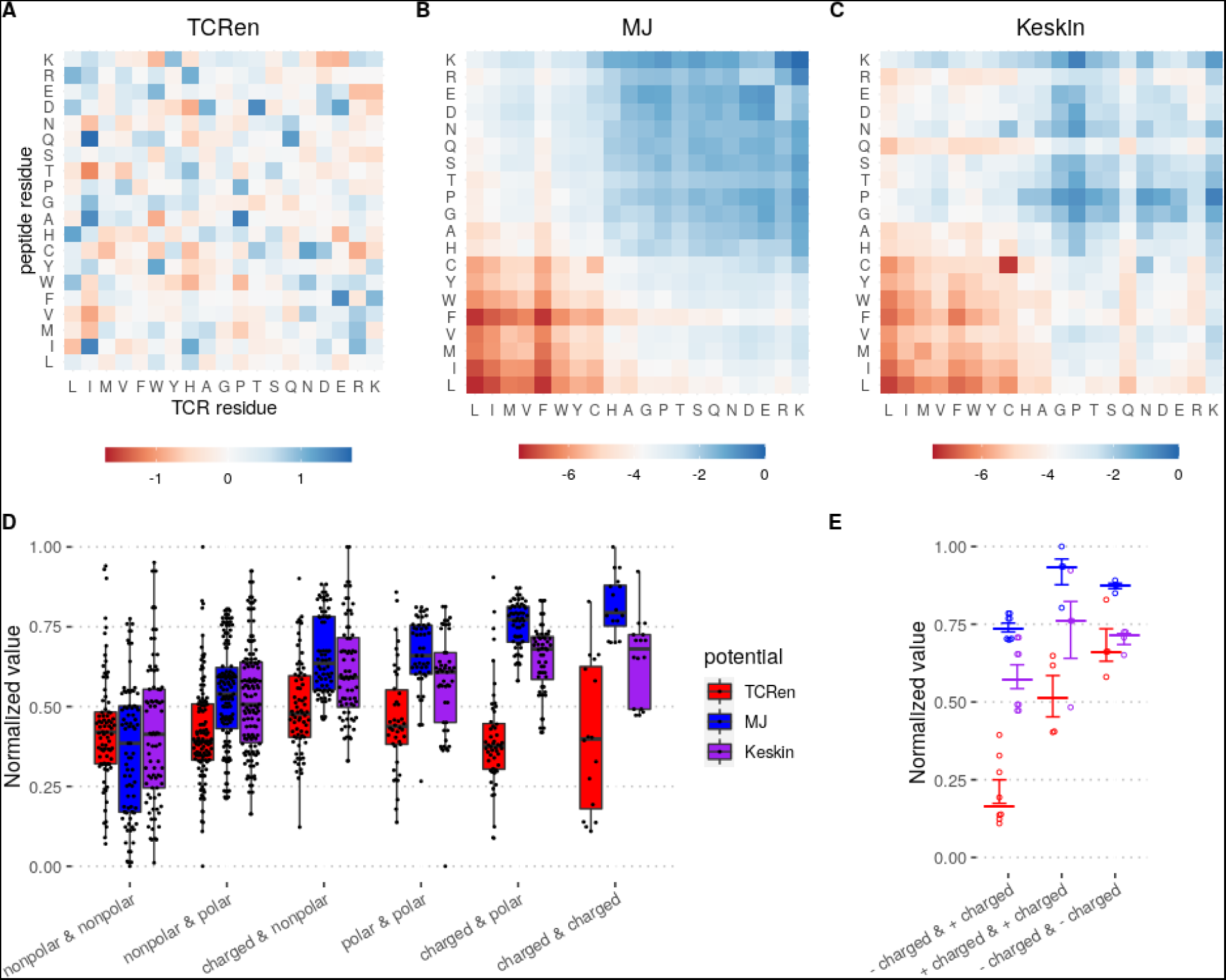
Comparison of TCRen, Miyazawa-Jernigan (MJ) and Keskin potentials. **a.** TCRen potential. **B.** MJ potential. **C.** Keskin potential. Color of each cell in **A.-C.** represents the estimated value of interaction energy between the residues denoted in the axes when they are in contact. **D-E.** Comparison of TCRen, MJ and Keskin values grouped by properties of the contacting amino acids. Values were normalized as: 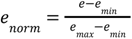, when *e_min_* and *e_max_* are the minimal and maximal values among entries of the given potential. Groups in **D.**: “nonpolar” — leucine, isoleucine, valine, methionine, phenylalanine, tryptophan, alanine, glycine, proline; “polar” — serine, threonine, asparagine, glutamine, histidine, cysteine, tyrosine; “charged” — arginine, lysine, aspartic acid, glutamic acid). In **E.** the group of “charged & charged” is decomposed to “+ charged” (arginine, lysine) and “− charged” (aspartic acid, glutamic acid).

**Supplementary Figure 5.**
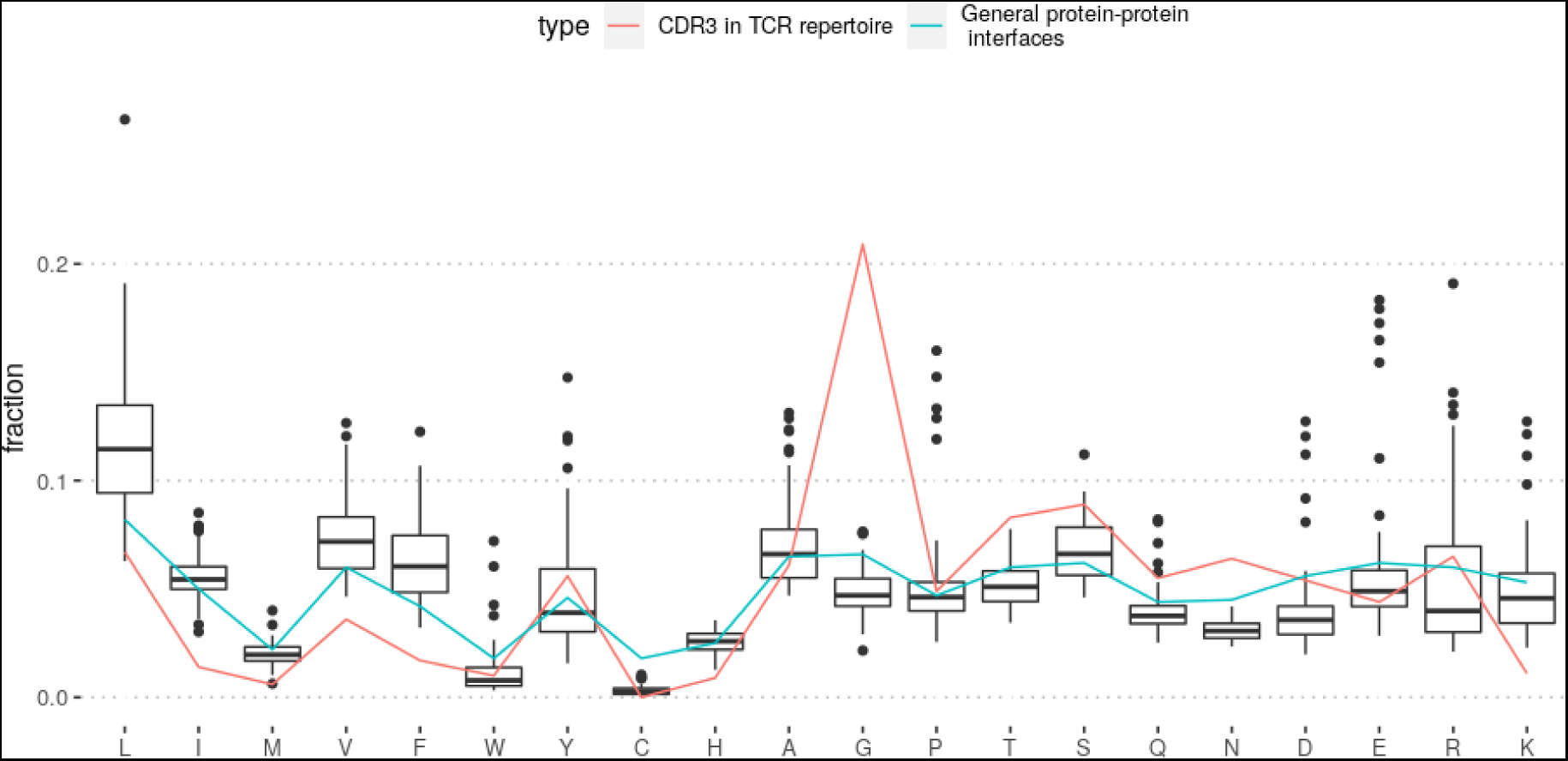
Amino acid composition of CDR3 regions of TCRs, MHC-presented peptides and general protein-protein interfaces. Boxplot shows distribution of fractions of amino acids in ligands for different HLA alleles (only HLA-A, HLA-B and HLA-C alleles, for which more than 1000 unique ligands are available in IEDB, are included, n = 54). Red and blue lines correspond to amino acid composition of CDR3 regions in TCR repertoire and general protein-protein interfaces, respectively. Note that CDR3 regions of TCRs are depleted in hydrophobic and enriched in polar amino acids compared to immunopeptidome and general protein-protein interfaces. Amino acid composition of the central parts of CDR3 regions in TCR repertoires was calculated from data by Emerson *et al.*^27^ and amino acid composition of MHC-presented peptides - from data of The Immune Epitope Database (IEDB)^22^.

**Supplementary Figure 6.**
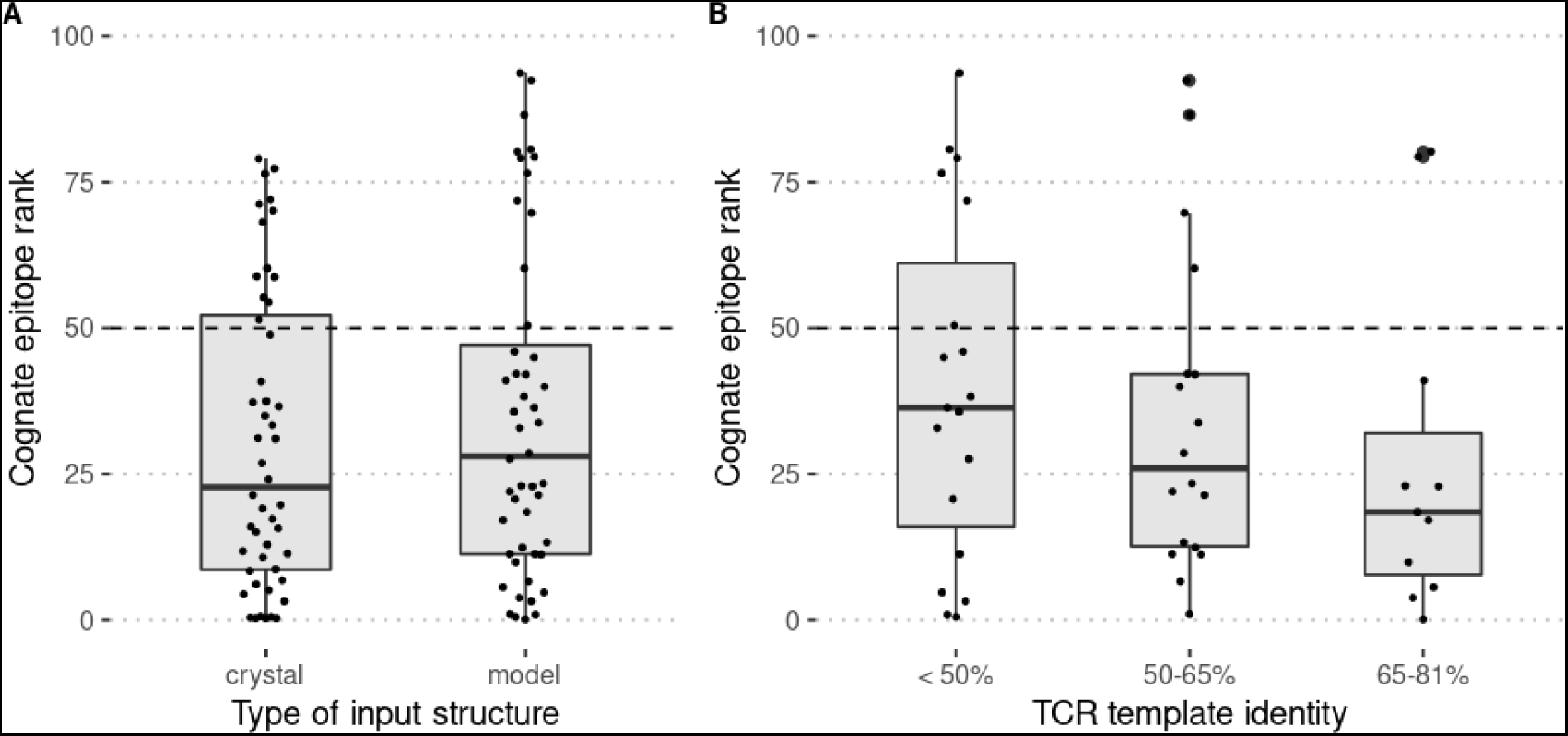
Performance of TCRen when homology models are used as input. **a.** Comparison of TCRen performance (cognate epitope rank) in distinction between cognate and unrelated peptides in the settings when either crystal structures or homology models are used as input. Homology models were obtained using TCRpMHCmodels tool; templates were manually checked to ensure that sequence identity of TCR structures used for modeling was less than 81% compared to target ones. The analysis was performed for a subset of recently published TCR-peptide-MHC structures, which are not included in TCRpMHCmodels template database. **B.** Relationship between performance TCRen (cognate epitope rank) and sequence similarity (identity) of the target TCR to the most similar one in the TCRpMHCmodels template database.

**Supplementary Figure 7.**
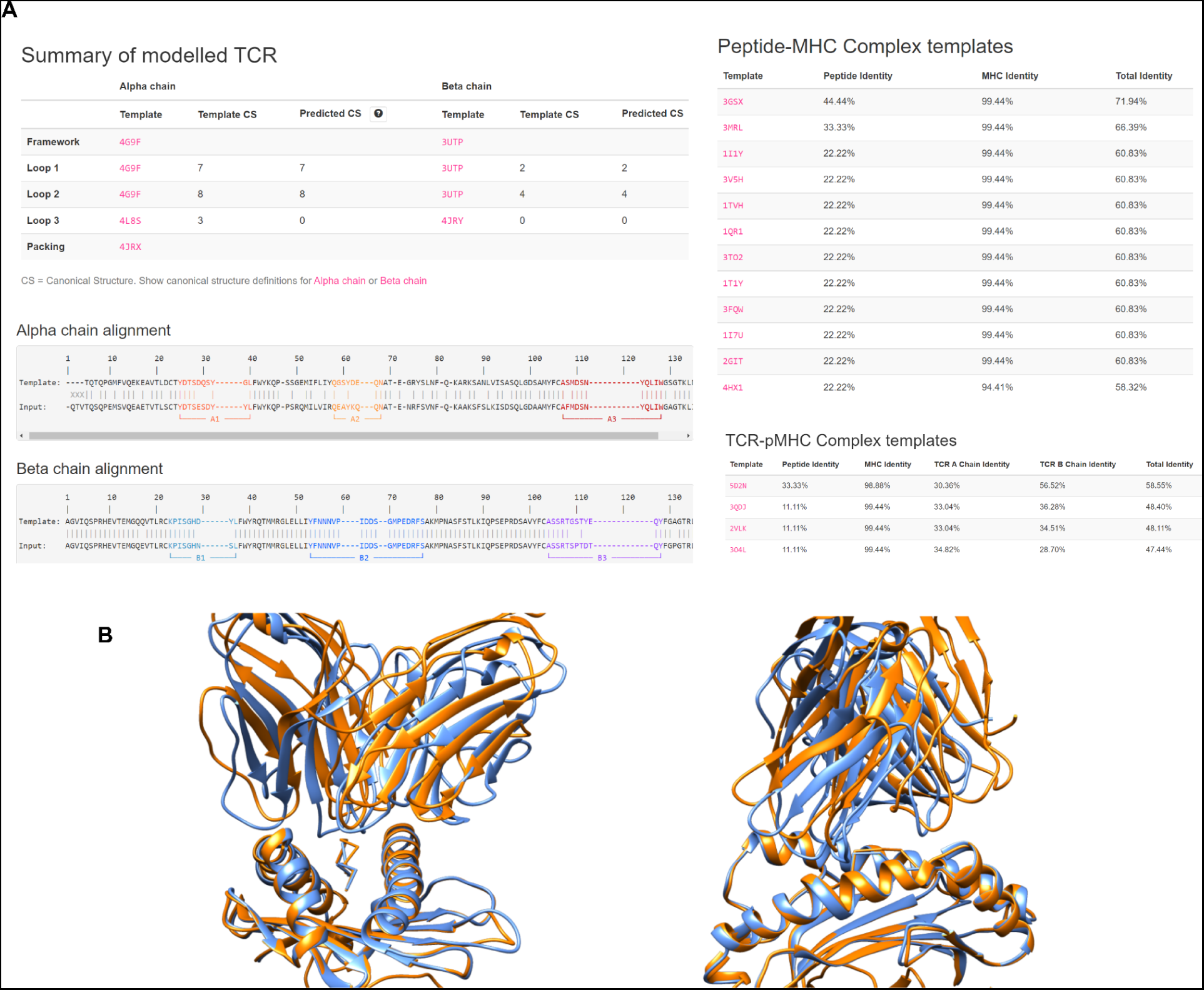
Homology modeling of 302TIL TCR-pepide-MHC complex. **a.** Templates used by TCRpMHCmodels server for homology modeling of 302TIL-VVVVVVVVV-HLA-A*02:06 complex. TCRpMHCmodels server at first performs homology modeling of TCR and peptide-MHC complex in separate and then assembles the full model. Templates selected for each stage are shown. **B.** Superposition of 302TIL-VVVVVVVVV-HLA-A*02:06 homology model (blue) and 302TIL-KQWLVWLFL-HLA-A*02:06 crystal structure (PDB ID: 6UK4; orange).

**Supplementary Figure 8.**
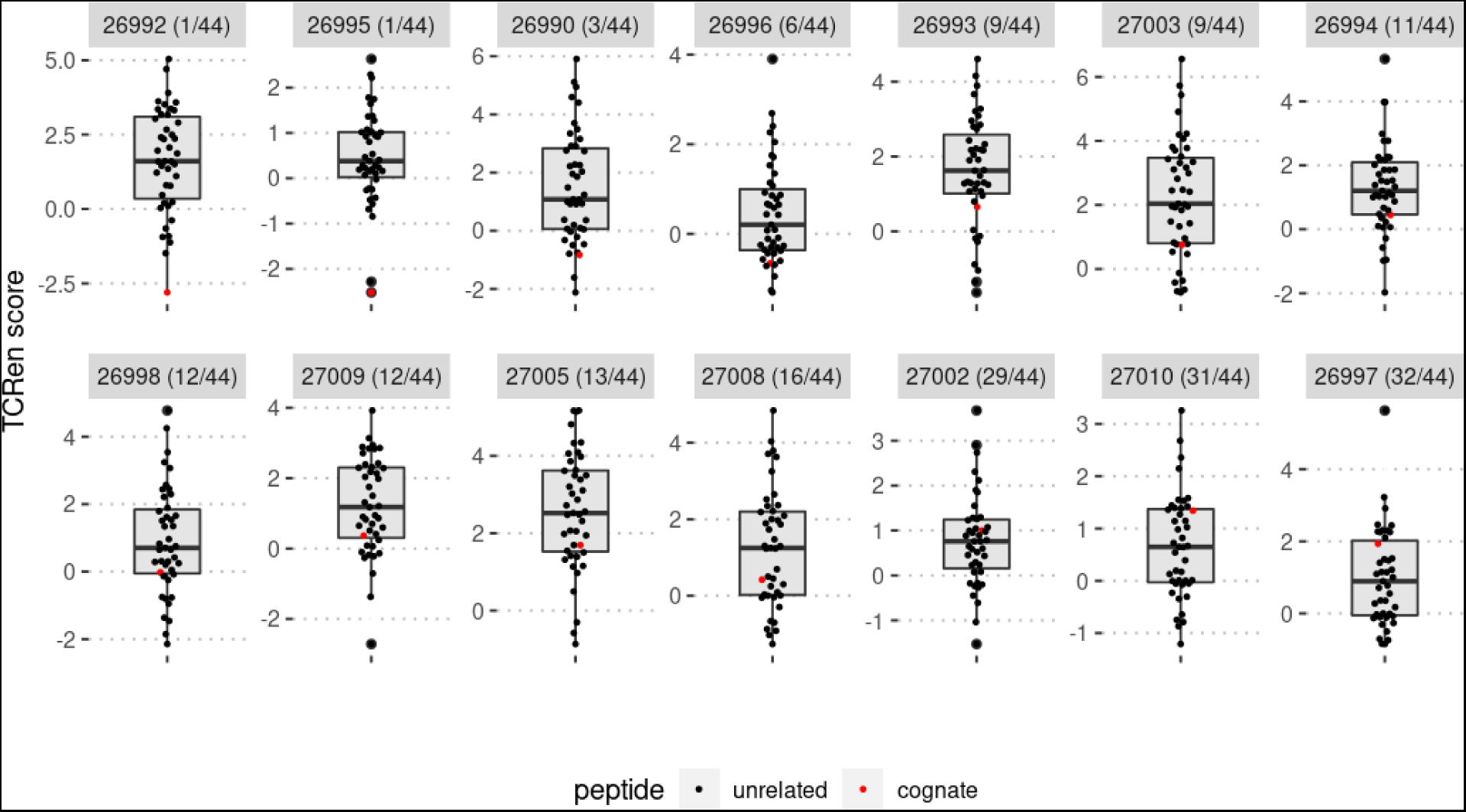
Identification of neoepitopes recognized by cancer-reactive T cells in SF3B1^mut^ uveal melanoma. TCR sequences of 14 tumor-reacting T cells and the list of 44 candidate neoepitopes predicted from *SF3B1*^mut^-modified intron–exon junctions were taken from the work of Bigot et al.^17^ For all 14 TCRs TCRen scores for all 44 candidate neoepitopes were calculated using the pipeline from **Figure 1b**. Distribution of TCRen scores of candidate neoepitopes for each TCR is shown, the cognate neoepitope is colored in red. The number in the header corresponds to “complex.id” of the TCR in the VDJdb database; the rank of the cognate epitope among 44 candidates is indicated in the brackets.

**Supplementary Figure 9.**
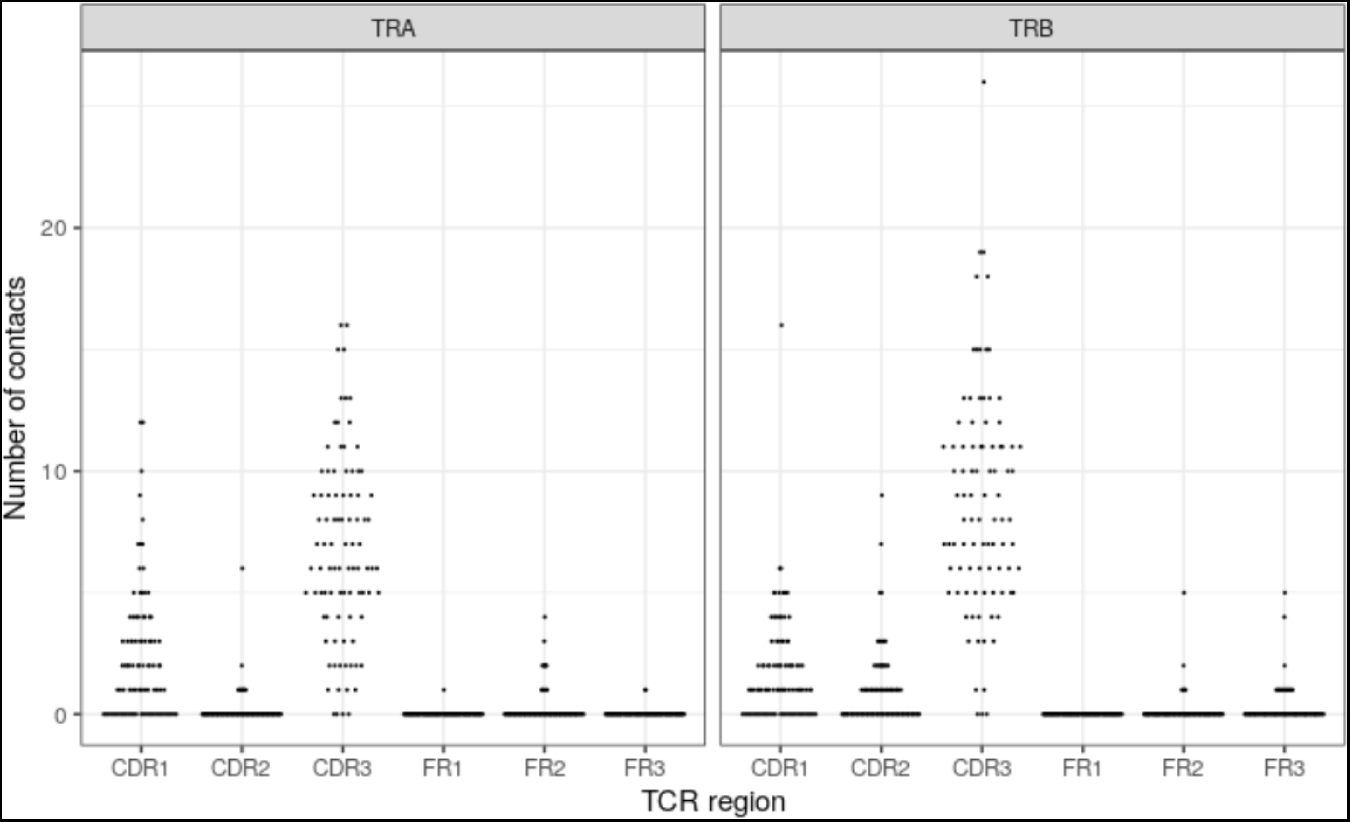
Distribution of the number of contacts of TCR complementarity determining (CDR) and framework (FR) regions with peptide in TCR-pMHC crystal structures of the non-redundant set used for TCRen derivation. Each point corresponds to the number of residue contacts between the given TCR region and peptide in a single structure from the non-redundant set.

**Supplementary Figure 10.**
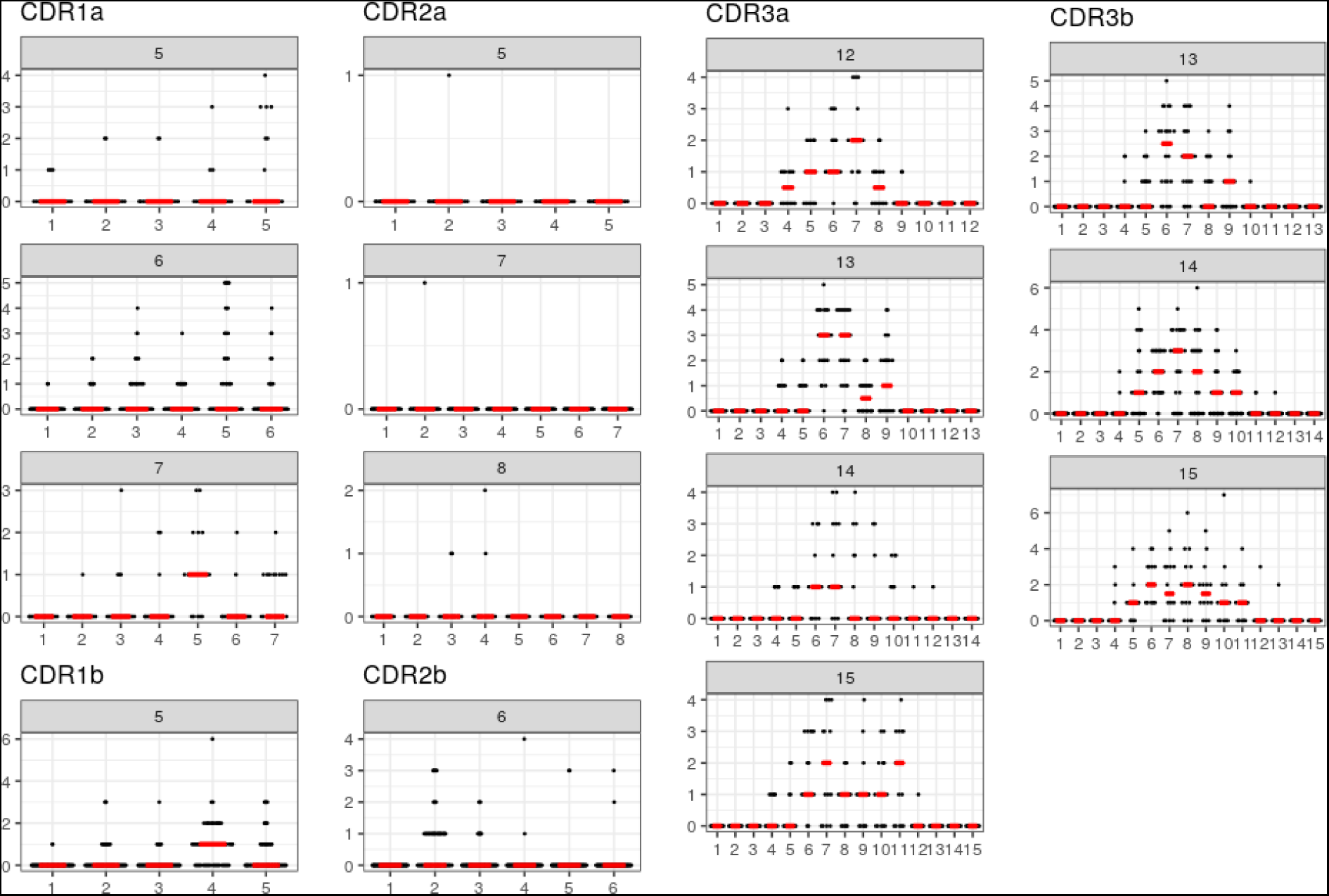
Relation between the residue position in CDR1/2/3 and the number of its contacts with the peptide. Number of contacts that residues in different positions of CDR1, CDR2 and CDR3 form with peptides in structures from the non-redundant set of TCR-peptide-MHC complexes. X axis value shows the residue position in CDR, Y axis - the number of contacts between the corresponding residue and the peptide. Red lines mark median values. Note that median values are 0 for almost all residues in CDR1 and CDR2. Each point corresponds to the number of contacts the residue in the given position in the given TCR forms with the peptide in a single structure from the non-redundant set. The distributions are shown only for the most values of CDR length, for which at least 10 structures in the non-redundant dataset exist with the same length of the given CDR.

**Supplementary Figure 11.**
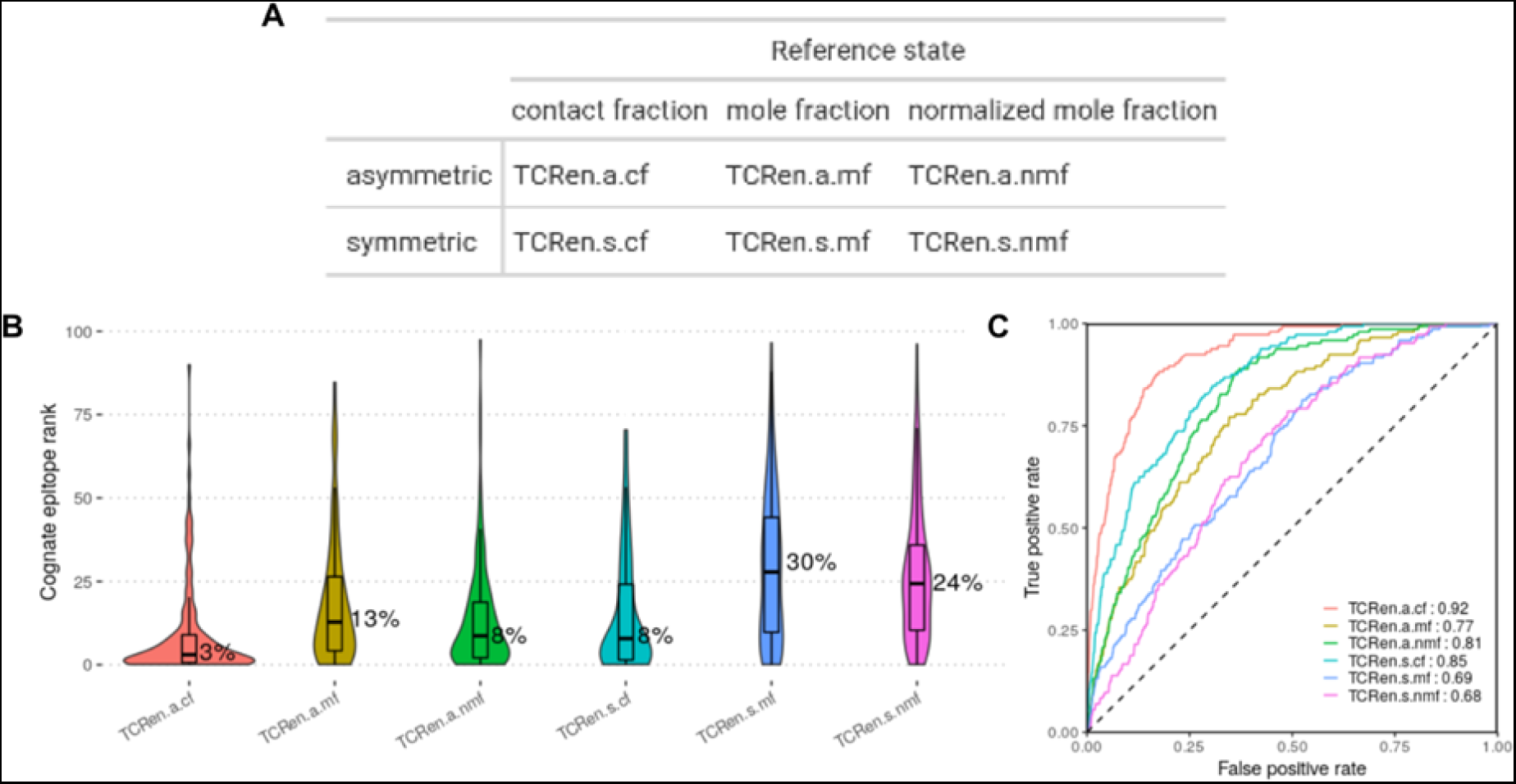
Comparison of performance of different versions of TCRen. **a.** Summary of notations used to refer to different versions of TCRen potential. All versions are called as TCRen.X.Y, when X is either “s” for symmetric or “a” for asymmetric forms, Y is either “cf” for contact-fraction or “mf” for mole-fraction or “nmf” for normalized mole-fraction reference states. **B.** Distributions of cognate epitope ranks for different versions of TCRen in cognate/unrelated peptides benchmark (benchmark setup is the same as in **Figure 2A**, see **Methods** for the details). Median value for each potential is marked. **C.** ROC-curves for distinction between all the complexes with cognate epitopes and all the complexes with unrelated peptides using different versions of TCRen.

**Supplementary Figure 12.**
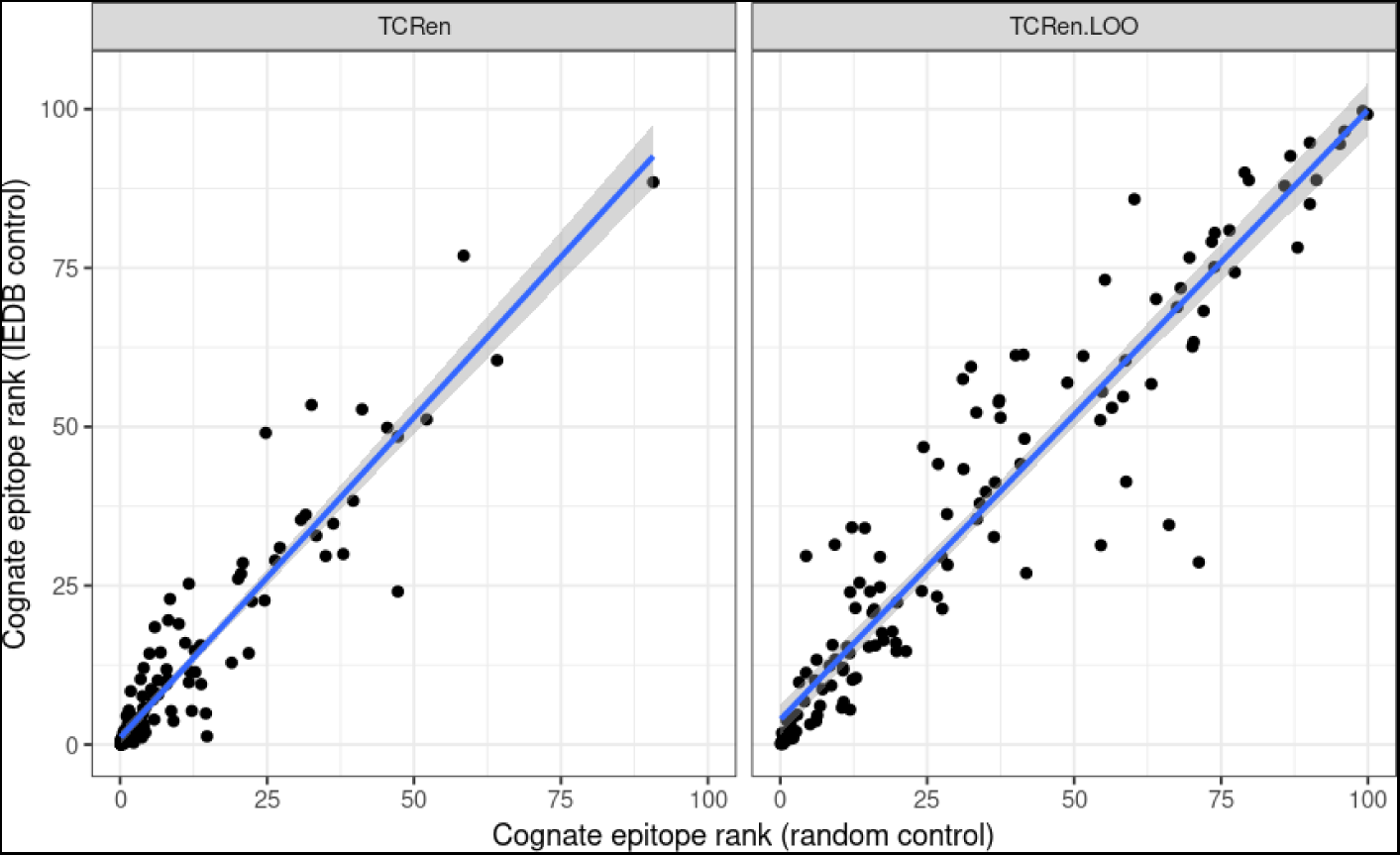
Absence of impact of strategy for choice of control peptides set on TCRen performance. Comparison of cognate epitope ranks in benchmark shown in **Figure 2a**, obtained when using different sets of unrelated peptides: either random mismatched peptides (x axis) or peptides with confirmed binding to the same MHC from IEDB (y axis).

**Supplementary Figure 13.**
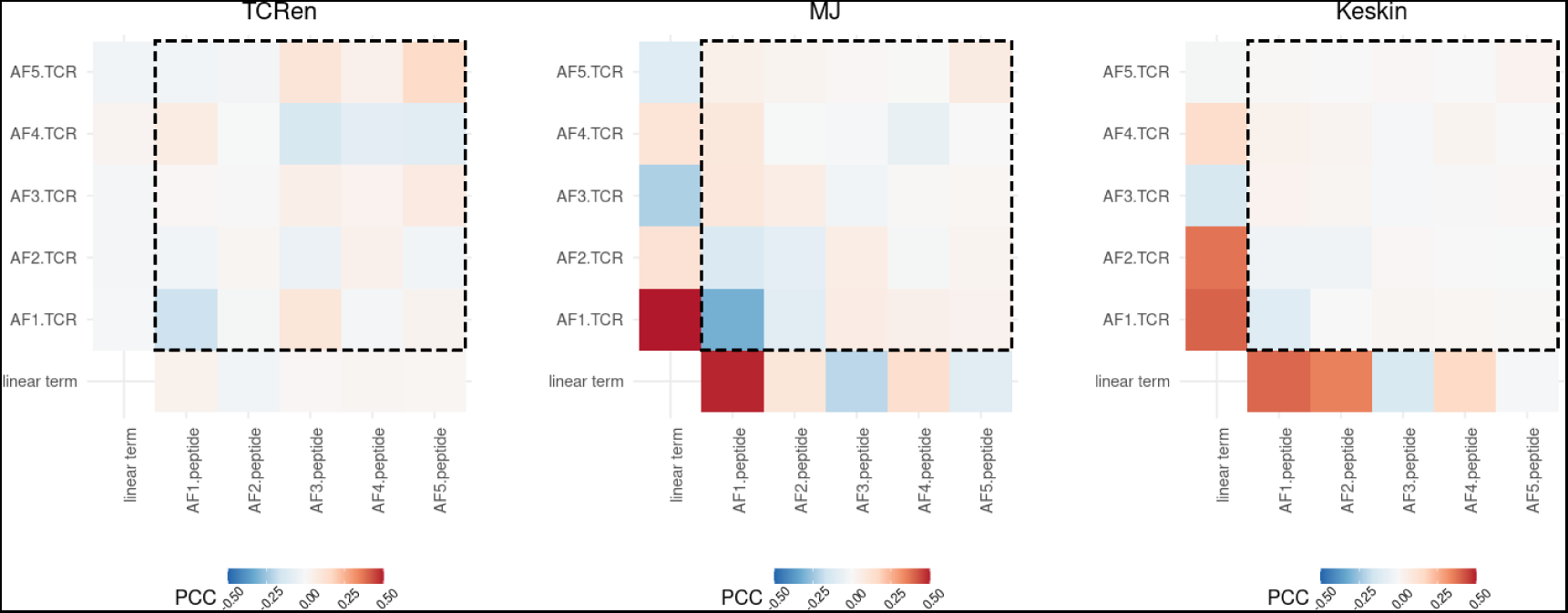
Correlation of TCRen, MJ and Keskin entries with Atchley factors (AFs) of contacting amino acids. Suffixes “.TCR” and “.peptide” denote residues from TCR and peptide, respectively. The first column corresponds to correlation coefficients of matrix entries with AFs of TCR residues, the last row — to correlation coefficients of matrix entries with AFs of peptide residues, and all other cells (marked with dashed rectangles) — to correlation coefficients between matrix entries and products of AFs, the first of which referred to residue from TCR and the second — to the residue from peptide. MJ and Keskin entries better correlate with properties of individual amino acids, while TCRen — with products of AFs of contacting amino acids. AF1: polarity/hydrophobicity, AF2: secondary structure preferences, AF3: molecular volume, AF4: codon diversity, AF5: electrostatic charge (see also **Supplementary Table 3**).

**Supplementary Figure 14.**
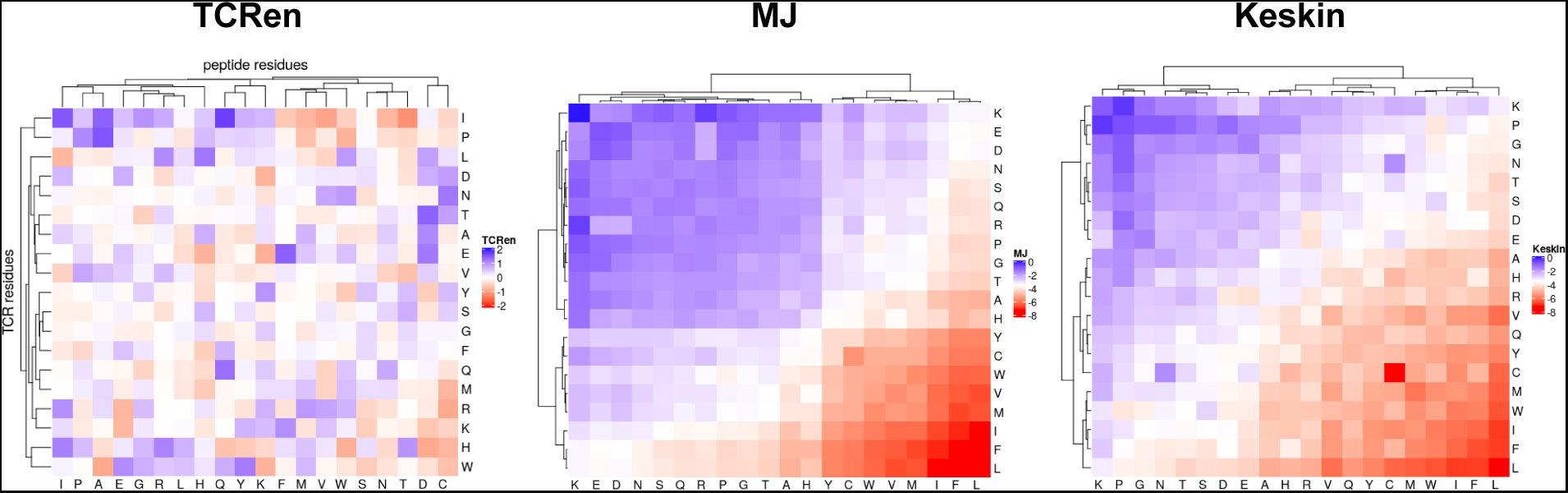
Clustering of TCRen, MJ and Keskin matrices. For TCRen matrix axis X corresponds to peptide residues, axis Y corresponds to TCR residues. Hierarchical clustering was performed based on the euclidean distance using stats::hclust R function.

**Supplementary Figure 15.**
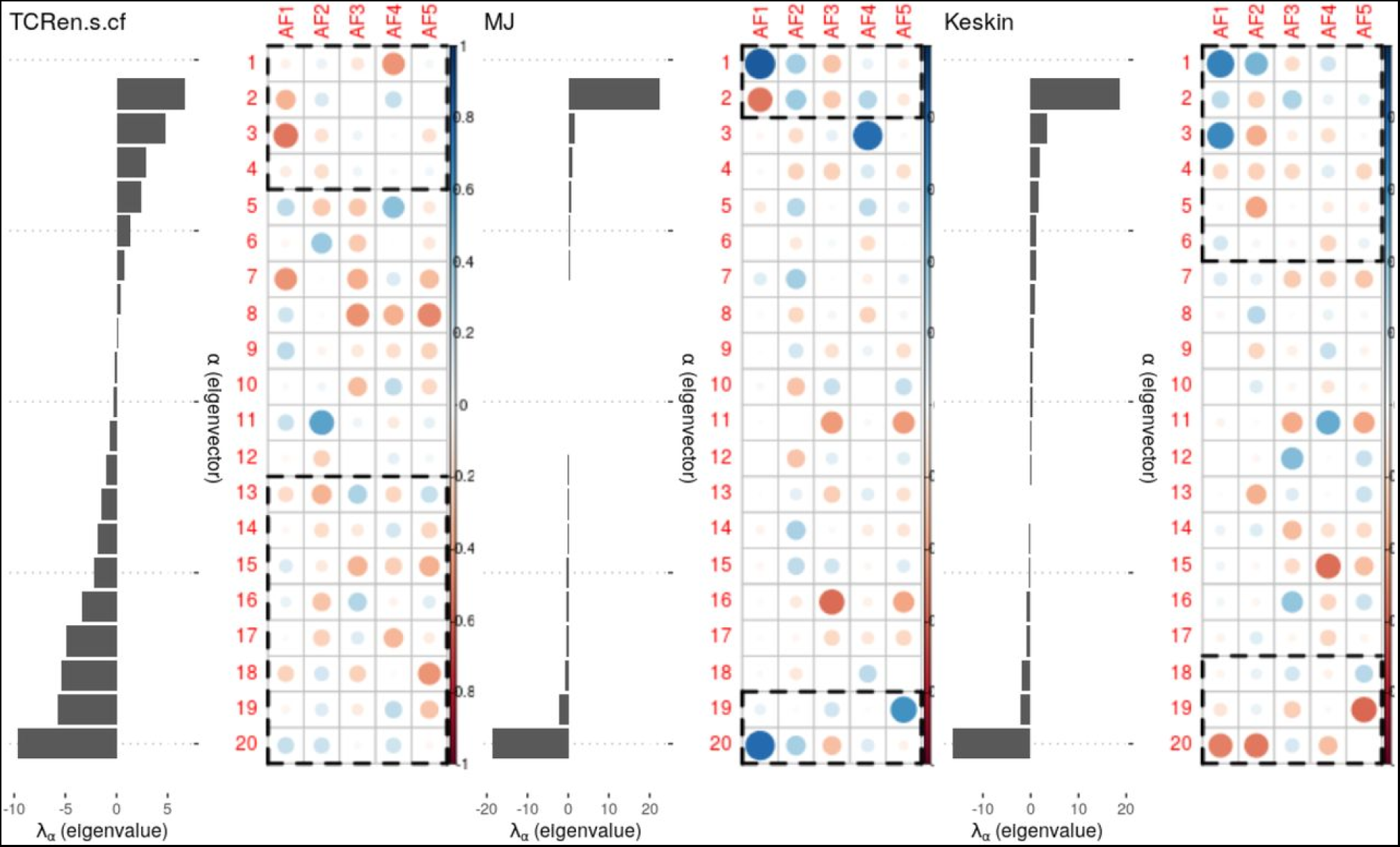
Eigenvalue decomposition for matrices of symmetrized TCRen, MJ and Keskin potentials. For each potential the left half of the figure represents eigenvalues, the right half shows correlation coefficients between corresponding eigenvectors and AFs of residues. For each matrix, eigenvectors which cumulatively explain 90% of variance are marked with dashed rectangles.

**Supplementary Figure 16.**
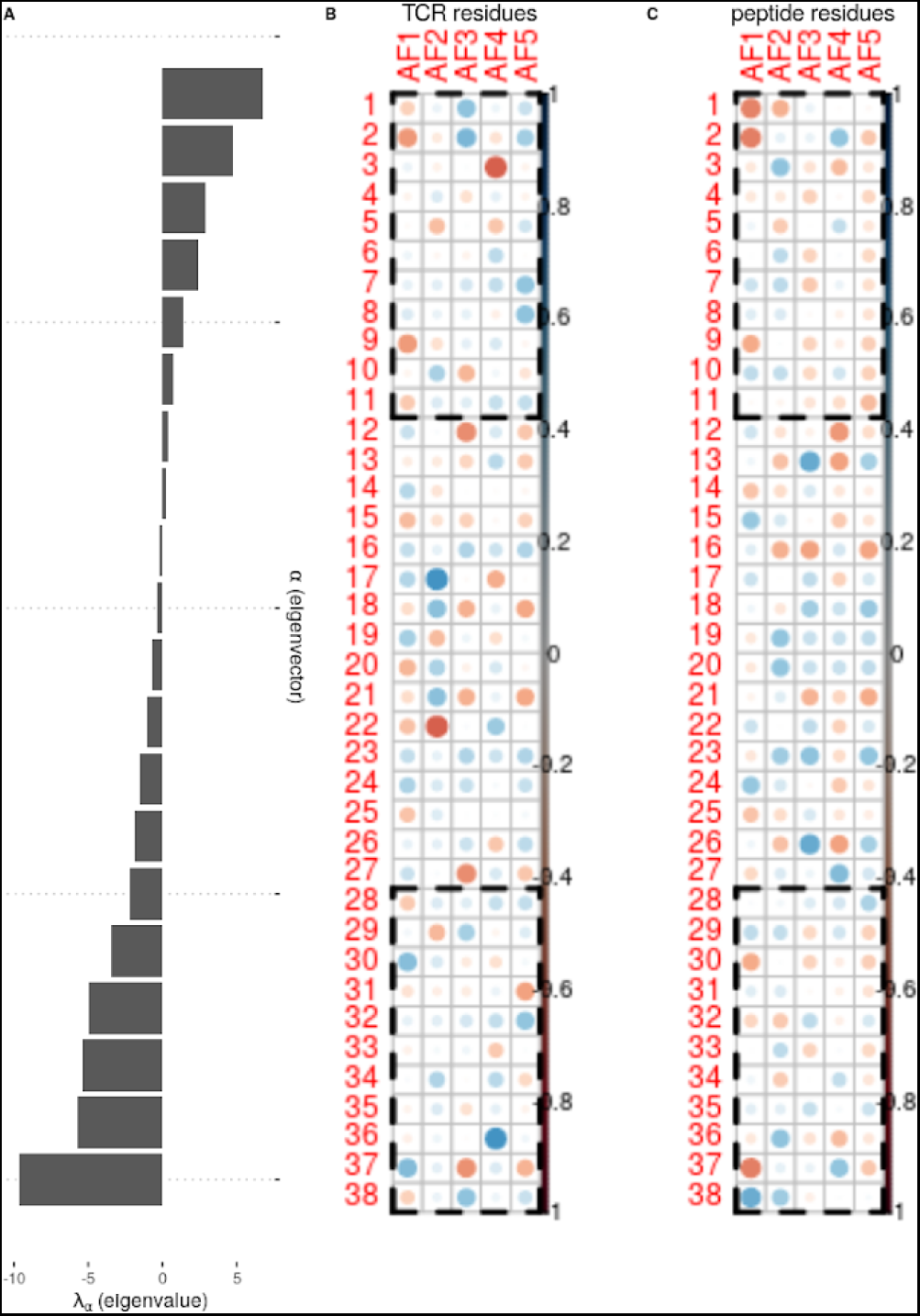
A. The eigenvalue spectrum for the TCRen matrix. **B-C.** Correlation of TCRen matrix eigenvectors with AF of TCR (**B**) and peptide (**C**) residues.

### Supplementary Tables

**Supplementary Table 1.**
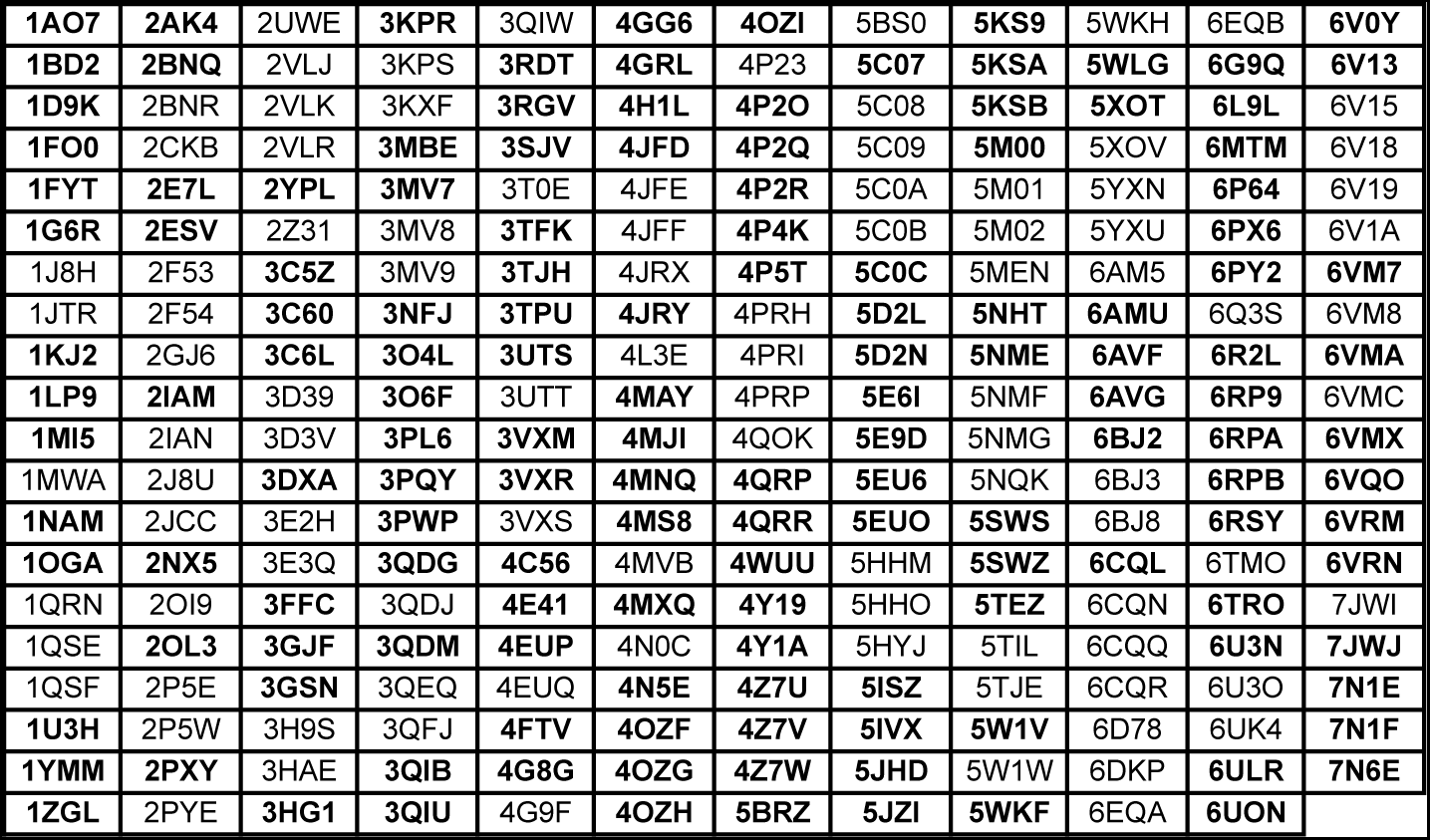
PDB IDs of all crystal structures of TCR-peptide-MHC complexes available at PDB. Structures included in the non-redundant set (**Supplementary Table 2**) are marked in bold.

**Supplementary Table 2.** Non-redundant set of crystal structures of TCR-peptide-MHC complexes from PDB used in the work for derivation of TCRen potential.

| PDB ID | Species | MHC class | peptide | CDR3a | CDR3b |
| --- | --- | --- | --- | --- | --- |
| 1AO7 | Human | MHCI | LLFGYPVYV | CAVTTDSWGKLQF | CASRPGLAGGRPEQYF |
| 1BD2 | Human | MHCI | LLFGYPVYV | CAAMEGAQKLVF | CASSYPGGGFYEQYF |
| 1D9K | Mouse | MHCII | GNSHRGAIEWEGIESG | CAATGSFNKLTf | CASGGQGRAEQFF |
| 1FO0 | Mouse | MHCI | INFDNTI | CAMRGDYGGSGNKLIF | CTCSADRVGNTLYF |
| 1FYT | Human | MHCII | PKYVKQNTLKLAT | CAVSESPFGNEKLTf | CASSSTGLPYGYTF |
| 1G6R | Mouse | MHCI | SIYRYGL | CAVSGFASALTf | CASGGGGTLYF |
| 1KJ2 | Mouse | MHCI | KVITFIDL | CAARYQGGRALIF | CTCSAAPDWGASAEITLYF |
| 1LP9 | Human | MHCI | ALWGFFPVL | CALFLASSSFskLVF | CASSDWVSYEQYF |
| 1MI5 | Human | MHCI | FLRGRAYGL | CILPLAGGTSYGKLTf | CASSLGQAYEQYF |
| 1NAM | Mouse | MHCI | RGYVYQGL | CAMRGDYGGSGNKLIF | CTCSADRVGNTLYF |
| 1OGA | Human | MHCI | GILGFVFTL | CAGAGSQGNLIF | CASSRSRSEYQYF |
| 1U3H | Mouse | MHCII | SRGGASQYRPSQ | CAASANSgTYQRF | CASGDAGGGYEYQF |
| 1YMM | Human | MHCII | ENPVVHFFKNIVTP | CATDTTSGTYKYIF | CSARDLTSGANNEQFF |
| 1ZGL | Human | MHCII | VHFFKNIVTPRTPG | CALSGGDSsyKLIF | CASSLADRVNTEAFF |
| 2AK4 | Human | MHCI | LPEPLPQGQLTAY | CALSGFYNTDKLIF | CASPGLAGEYEYQF |
| 2BNQ | Human | MHCI | SLLMWITQV | CAVRPTSGGSYIPTf | CASSYVGNtGELFF |
| 2E7L | Mouse | MHCI | QLSPFPFDL | CAVSHQGRYLTF | CASGGGGTLYF |
| 2ESV | Human | MHCI | VMAPRTLIL | CIVVRSSNTGKLIF | CASSQDRDTQYF |
| 2IAM | Human | MHCII | GELIGILNAAKVPAD | CAALIQAQKLVF | CASrYHGTGYF |
| 2NX5 | Human | MHCI | EPLPQGQLTAY | CAVQASGGSYIPTf | CATGTGDSNQPQHf |
| 2OL3 | Mouse | MHCI | SQYYNSL | CAMRGDYGGSGNKLIF | CTCSADRVGNTLYF |
| 2PXY | Mouse | MHCII | RGGASQYRPSQ | CALSENYGNEKITf | CASGDASGAETLYF |
| 2YPL | Human | MHCI | KAFSPEVIPMF | CAVSGGYQKVTF | CASrGSYGYTF |
| 3C5Z | Mouse | MHCII | FEAQKAKANKAVDG | CALVISNTNKVVF | CASIDSSGNTLYF |
| 3C60 | Mouse | MHCII | FEAQKAKANKAVD | CAANSgTYQRF | CASGDFWGDtLYF |
| 3C6L | Mouse | MHCII | FEAQKAKANKAV | CAASDNRIFF | CASGDAWGYEYQF |
| 3DXA | Human | MHCI | EENLLDFVRF | CIVWGGYQKVTF | CASRYRDDSYNEQFF |
| 3FFC | Human | MHCI | FLRGRAYGL | CAMREDTGNQFYF | CASSFTWTSGGATDTQYF |
| 3GJF | Human | MHCI | SLLMWITQV | CAGELLPYGMDVW | CWSFAGSYVVF |
| 3GSN | Human | MHCI | NLVPMTATV | CARNTGNQFYF | CASSPVTGGIYGTYF |
| 3HG1 | Human | MHCI | ELAGILTV | CAVNvAGKSTf | CAWSETGLGTGELFF |
| 3KPR | Human | MHCI | EEYLKAWTF | CILPLAGGTSYGKLTf | CASSLGQAYEQYF |
| 3MBE | Mouse | MHCII | GAMKRHGLDNYRGYSLG | CAEDGGSGNKLIF | CASSWDRAgNTLYF |
| 3MV7 | Human | MHCI | HPVGEADYFEY | CAVQDLGTSGSRLTf | CASSARSgELFF |
| 3NFJ | Human | MHCI | RYPLTFGWCF | CALGELARSGGYQKVTF | CAWSVSVGAGVPTIYF |
| 3O4L | Human | MHCI | GLCTLVAML | CAEDNNARLMF | CSARDGTNGYTF |
| 3O6F | Human | MHCII | FSWGAEGQRPGF | CTVYGGATNKLIF | CSARGGSYNsPLHF |
| 3PL6 | Human | MHCII | NPVVHFFKNIVTPR | CAASSFGNEKLTf | CATSALGDTQYF |
| 3PQY | Mouse | MHCI | SSLENFRAYV | CILSGGSNYKLTf | CASSFGREQYF |
| 3PWP | Human | MHCI | LGYGfVNYI | CAVTTDSWGKLQF | CASRPGLAGGRPEQYF |
| 3QDG | Human | MHCI | ELAGILTV | CAVNfGGGKLIF | CASSLSFGTEAFF |
| 3QDM | Human | MHCI | ELAGILTV | CAGGTGNQFYF | CAISEVGVGGPQHf |
| 3QIB | Mouse | MHCII | ADLIAYLKQATKG | CAALRATGGNNKLTf | CASSLNWSQDTQYF |
| 3QIU | Mouse | MHCII | ADLIAYLKQATKG | CAAEPSGGKLVF | CASSLNANSdYTF |
| 3RDT | Mouse | MHCII | FEAQKAKANKAVD | CAASKGADRLTF | CASGDFWGDtLYF |
| 3RGV | Mouse | MHCI | WIYVYRPMGCGGS | CAANSgTYQRF | CASGDFWGDtLYF |
| 3SJV | Human | MHCI | FLRGRAYGL | CVVRAGKLIF | CASGQGNfDIQYF |
| 3TFK | Mouse | MHCI | QLSDVPMDL | CAVSAKGTGSKLSF | CASSDAPGQLYF |
| 3TJH | Mouse | MHCI | SPLDSLWWI | CAVSAKGTGSKLSF | CASSDAPGQLYF |
| 3TPU | Mouse | MHCI | FLSPFWFDI | CAVSAKGTGSKLSF | CASSDAPGQLYF |
| 3UTS | Human | MHCI | ALWGPDAAA | CAMRGDSsyKLIF | CASSLWEKLAKNIQYF |
| 3VXM | Human | MHCI | RFPLTFGWCF | CAVGAPSGAGSYQLTF | CASSPTSGIYEYQYF |
| 3VXR | Human | MHCI | RYPLTFGWCF | CAVRMDSSyKLIF | CASSSWDTGELFF |
| 4E41 | Human | MHCII | GELIGILNAAKVPAD | CAVDRGSTLGRLYF | CASSQIRETQYF |
| 4EUP | Human | MHCI | ALGIGILTV | CAVSGGGADGLTF | CASSFLGTGVEYQYF |
| 4FTV | Human | MHCI | LLFGYPVYV | CAVTTDSWGKLQF | CASRPGLMSAQPEQYF |
| 4G8G | Human | MHCI | KRWIILGLNK | CAMRDLRDnFNKFYF | CASREGLGGTEAFF |
| 4GG6 | Human | MHCII | SGEGSFQPSQENP | CILRDGRGGADGLTF | CASSVAVSAGTYEQYF |
| 4GRL | Human | MHCII | RLLMLFAKDVVSRN | CAASSFGNEKLTf | CATSALGDTQYF |
| 4H1L | Human | MHCII | QHIRCNIPIKRISA | CAVGASGNTGKLIF | CASSLRDGYTGELFF |
| 4JFD | Human | MHCI | ELAAIGILTV | CAVNDGGRLTF | CAWSETGLMGGWQF |
| 4JRY | Human | MHCI | LPEPLPQQQLTAY | CAVGGSNSYQLIW | CASSRTGSTYEQYF |
| 4MAY | Human | MHCII | QLVHFVRDFAQL | CAASSFGNEKLTf | CATSALGDTQYF |
| 4MJI | Human | MHCI | TAFTIPSI | CATDDDSARQLTF | CASSLTGGGELFF |
| 4MNQ | Human | MHCI | ILAKFLHWL | CAVDSATALPYGYIF | CASSYQGTEAFF |
| 4MS8 | Mouse | MHCI | SPAEAGFFL | CAVSAKGTGSKLSF | CASSDAPGQLYF |
| 4MXQ | Mouse | MHCI | SPAPRPLDL | CAVSAKGTGSKLSF | CASSDAPGQLYF |
| 4N5E | Mouse | MHCI | VPYMAEFGM | CAVSAKGTGSKLSF | CASSDAPGQLYF |
| 4OZF | Human | MHCII | APQPELPYPQPGS | CIAFQGAQKLVF | CASSFRALAADTQYF |
| 4OZG | Human | MHCII | APQPELPYPQPG | CIVLGGADLTF | CASSFRFTDTQYF |
| 4OZH | Human | MHCII | APQPELPYPQPGS | CIVWGGATNKLIF | CASSVRSTDTQYF |
| 4OZI | Human | MHCII | QFPQPPELPYP | CLVGDGGSFSGGYNKLIF | CSAGVGGQETQYF |
| 4P2O | Mouse | MHCII | PADPLAFFSSAIKGGGSLV | CAALRATGGNNKLTF | CASSLNWSQDTQYF |
| 4P2Q | Mouse | MHCII | ADGLAYFRSSFQGG | CAAEASNTNKVVF | CASSLNNANSDYTF |
| 4P2R | Mouse | MHCII | ANGVAFFLTPFKA | CAAEASNTNKVVF | CASSLNNANSDYTF |
| 4P4K | Human | MHCII | QAFWIDLFETIG | CALSLYSGAGSYQLTF | CASSLAQGGETQYF |
| 4P5T | Mouse | MHCII | FEAQKAKANKAVD | CAASRDGQKLVF | CASGDFWGDTLF |
| 4QRP | Human | MHCI | HSKKKCDL | CALSDPVNDMRF | CASSLRGRGDQPQHF |
| 4QRR | Human | MHCI | IPSINVHHY | CALGELAGAGGTSYGKLTf | CASSLEGYYNEQFF |
| 4WUU | Human | MHCI | RMFPNAPYL | CARVQYSGYYDWFDPW | CAAWDDSLNGWVF |
| 4Y19 | Human | MHCII | QPLALEGSLQKRG | CAASVYAGGTSYGKLTf | CASRPDRDNEQFF |
| 4Y1A | Human | MHCII | LQPLALEGSLQKRG | CAASSAGGTSYGKLTf | CASRPDRPVDTQYF |
| 4Z7U | Human | MHCII | APSGEGSFQPSQENPQ | CILRDRSNQFYF | CASSTTPGTGTETQYF |
| 4Z7V | Human | MHCII | SGEGSFQPSQENP | CILRDSRAQKLVF | CASSAGTSGEYEQYF |
| 4Z7W | Human | MHCII | PSGEGSFQPSQENPQ | CAVGETGANNLFF | CASSEARRYNEQFF |
| 5BRZ | Human | MHCI | EVDPIGHL | CAVRPGGAGPFFVVF | CASSFNMATGQYF |
| 5C07 | Human | MHCI | YQFGPDFPIA | CAMRGDSSYKLIF | CASSLWEKLAKNIQYF |
| 5C0C | Human | MHCI | RQFGPDWIVA | CAMRGDSSYKLIF | CASSLWEKLAKNIQYF |
| 5D2L | Human | MHCI | NLVPMTATV | CAFITGNQFYF | CASSQTLWETQYF |
| 5D2N | Human | MHCI | NLVPMTATV | CILDNNNDMRF | CASSLAPGTTNEKLFF |
| 5E6I | Human | MHCI | GILGFVFTL | CAGPGGSSNTGKLIF | CASSLIYPGELFF |
| 5E9D | Human | MHCI | ELAGIGILTV | CAVTKYSWGKLQF | CASRPGWMMAGGVELYF |
| 5EU6 | Human | MHCI | YLEPGPVTV | CAVLSSGGSNYKLTF | CASSFIGGTDQYF |
| 5EUO | Human | MHCI | GILGFVFTL | CAGAIGPSNTGKLIF | CASSIRSSYEQYF |
| 5ISZ | Human | MHCI | GILGFVFTL | CAFDTNAGKSTF | CASSIFGQREQYF |
| 5IVX | Mouse | MHCI | RGPGRAFVTI | CAASASFGDNSKLIW | CASSLGHTVEVFF |
| 5JHD | Human | MHCI | XGILGFVFTL | CAWGVNAGGTSYGKLTf | CASSIGVYGYTF |
| 5JZI | Human | MHCI | XKLVALGINAV | CAYGEDDKIIF | CASRRGPYEQYF |
| 5KS9 | Human | MHCII | APSGEGSFQPSQENPQ | CAVALNNNAGNMLTF | CASSVAPGSDTQYF |
| 5KSA | Human | MHCII | QPQQSFPEQEA | CAVQFMDSNYQLIW | CASSVAGTPSYEQYF |
| 5KSB | Human | MHCII | GPQQSFPEQEA | CAVQASGGSYIPTF | CASSNRGLGTDQYF |
| 5M00 | Mouse | MHCI | KAVANFATM | CAALYGNEKITF | CASSDAGGRNTLYF |
| 5NHT | Human | MHCI | ELAGIGILTV | CAVGGGADLTF | CASSQGLAGAGELFF |
| 5NME | Human | MHCI | SLYNTVATL | CAVRTNSGYALNF | CASSDTSVSYEQYF |
| 5SWS | Mouse | MHCI | ASNENMETM | CAASEGSGSWQLIF | CASSAGLDAEQYF |
| 5SWZ | Mouse | MHCI | ASNENMETM | CAASETSGSWQLIF | CASSRDLGRDTQYF |
| 5TEZ | Human | MHCI | GILGFVFTL | CAASFIQGAQKLVF | CASSLLGGWSEAFF |
| 5W1V | Human | MHCI | VMAPRTLIL | CAGQPLGGSNYKLTF | CASSANPGDSSNEKLFF |
| 5WKF | Human | MHCI | GTSGSPIVNR | CGLGDAGNMLTF | CASSLGQGLLYGYTF |
| 5WLG | Mouse | MHCI | SQLLNAKYL | CATVYAQGLTF | CASSDWGDTGQLYF |
| 5XOT | Human | MHCI | IPLTEEAEL | CALGEGGAQKLVF | CASRTRGGLIEQYF |
| 6AMU | Human | MHCI | MMWDRGLGMM | CAVNFGGKLI | CASSLSFGTEAFF |
| 6AVF | Human | MHCI | APRGPHGGAASGL | CLVGEILDNFNKFYF | CASSQRQEGDTQYF |
| 6AVG | Human | MHCI | APRGPHGGAASGL | CLVVDQKLVF | CASSGGHTGSNEQFF |
| 6BJ2 | Human | MHCI | IPLTEEAEL | CALSHNSGGSNYKLTF | CASSFRGKGTQYF |
| 6CQL | Human | MHCII | RFYKTLRAEQASQ | CAFKAAAGNKLTF | CASSRLAGGMDEQFF |
| 6G9Q | Mouse | MHCI | KAPYDYAPI | CAALYGNEKITF | CASSDAGGRNTLYF |
| 6L9L | Mouse | MHCI | SPSYAYHQF | CILQGTGSKLSF | CASSDGDYEQYF |
| 6MTM | Human | MHCI | FEDLRVLSF | CGTERSGGYQKVTF | CASSMSAMGTEAFF |
| 6P64 | Human | MHCI | KQWLVLFL | CAFMDSNYQLIW | CASSRTSPTDTQYF |
| 6PX6 | Human | MHCII | APFSEQEQPVLG | CAVHTGARLMF | CASSHGASTDTQYF |
| 6PY2 | Human | MHCII | APFSEQEQPVLG | CASPQGGSEKLVF | CASSSGGWGGGTEAFF |
| 6R2L | Human | MHCI | SLSKILDTV | CAVGNDWNTDKLIF | CASSPLDVSISSYNEQFF |
| 6RP9 | Human | MHCI | SLLMWITQV | CALTRGPGNQFYF | CASSSPGGVSTEAFF |
| 6RPA | Human | MHCI | SLLMWITQV | CAVRDINSAGASYQLTF | CSVGGSGGADTQYF |
| 6RPB | Human | MHCI | SLLMWITQV | CAVKSGGSYIPTF | CASSYLNKDSALDF |
| 6RSY | Human | MHCI | RMFPNAPYL | CIGGGTTSPTYKYIF | CASSLGFGKRDVMRF |
| 6TRO | Human | MHCI | GVYDGREHTV | CVVNHSGGSYIPTF | CASSFLMTSGDPYEQYF |
| 6U3N | Human | MHCII | APMPPELPYP | CAVGAGSNYQLIW | CASSLEGQGASEQFF |
| 6ULR | Human | MHCI | GADGVGKSA | CLVGDMQAGTALIF | CASSLGEGKRDGYTF |
| 6UON | Human | MHCI | GADGVGKSAL | CAAAMDSSYKLIF | CASSDPGTEAFF |
| 6V0Y | Human | MHCII | GGYAPAKAAAT | CALSDSGSFNKLTF | CASSLDWGGQNTLYF |
| 6V13 | Human | MHCII | GGYAPAKAAAT | CALSPSNTNKVVF | CASSLDWGVNTLYF |
| 6VM7 | Human | MHCI | ITDQVPFSV | CAVNARRNTPLVF | CASSITLSSYNEQFF |
| 6VMA | Human | MHCI | ITDQVPFSV | CAVNALLGNQFYF | CASSMGGTYEQYF |
| 6VMX | Human | MHCI | RPPIFIRRL | CAFGSSNTGKLIF | CASSQDLFTGGYTF |
| 6VQO | Human | MHCI | HMTEVVRHC | CAMSGLKEDSSYKLIF | CASSIQQGADTQYF |
| 6VRM | Human | MHCI | HMTEVVRHC | CVVQPGGYQKVT | CASSEGLWQVGDEQYF |
| 6VRN | Human | MHCI | HMTEVVRHC | CAFMGYSGAGSYQLTF | CAISELVTGDSPLHF |
| 7JWJ | Mouse | MHCI | ASNENMETM | CAAVTGNTGKLIF | CASSRGTHSNTEVFF |
| 7N1E | Human | MHCI | RLQSLQTYV | CALSGFNNAGNMLTF | CASSLGGAGGADTQYF |
| 7N1F | Human | MHCI | YLQPRTFLL | CAVNRDDKIIF | CASSPDIEQYF |
| 7N6E | Human | MHCI | YLQPRTFLL | CVVNRNNDMRF | CAGQVTNTGELFF |

**Supplementary Table 3.** Atchley factors.

| **Atchley factor** | **Property** | **Higher values** | **Lower values** |
| --- | --- | --- | --- |
| AF1 | Polarity/hydrophobicity | Hydrophilicity | Hydrophobicity |
| AF2 | Secondary structure | Bends, coils, turns | alpha-helical |
| AF3 | Molecular volume | Small size | Large size |
| AF4 | Codon diversity | Higher amino acid usage | Heat capacity, refractivity |
| AF5 | Electrostatic charge | Positively charged | Negatively charged |

### Supplementary Note 1

#### TCRen correlation with physicochemical properties of contacting amino acids

As TCRen was derived based on amino acid contact preferences in TCR:peptide interface, it should reflect the principles of TCR-peptide recognition. We performed the analysis of correlation between TCRen entries (**Supplementary Figure 4A**) and physicochemical properties of contacting amino acids to shed light on the specific nature of TCR-peptide interactions. For comparison we used the potential derived by Miyazawa and Jernigan^10^ (MJ; **Supplementary Figure 4B**), representing the rules governing protein folding, and the potential from the work of Keskin *et al.*^9^ (Keskin; **Supplementary Figure 4C**), reflecting preferences in general protein-protein interactions.

To investigate regularity patterns in TCRen, MJ and Keskin potentials we first grouped non-polar, polar and charged amino acids and compared values for contacts between two non-polar residues, one non-polar and one polar residue, one non-polar and one charged residue, etc. (**Supplementary Figure 4D**). MJ demonstrates clear clustering of non-polar, polar and charged residues (**Supplementary Figure 14**) with the best values for contacts of two non-polar residues and the worst values for contacts of two charged residues (**Supplementary Figure 4D**). Keskin follows in general the same patterns as MJ but with more favorable energy score values for contacts of oppositely charged residues (**Supplementary Figure 4E**) and contacts of glutamine and arginine with other amino acids (**Supplementary Figure 4C**). For TCRen some clustering of amino acids with similar properties (e.g. positively charged R and K, polar T, N and S, hydrophobic W, V, M, F) is observed (**Supplementary Figure 14**), however, generally it doesn’t have a simple regular pattern analogous to that of MJ and Keskin.

Further we examined correlation between TCRen, MJ and Keskin values and physicochemical properties of contacting residues represented as 5 Atchley factors (AF)^28^, which were derived from a multivariate analysis with dimension reduction of 494 properties of amino acids (**Supplementary Figure 13**). MJ and Keskin entries to a greater extent correlate with polarity/hydrophobicity AF1 of individual amino acids. On the opposite, TCRen correlates mostly with the products of AFs of contacting residues that implies that TCRen accounts for interaction of physicochemical properties of contacting amino acids.

#### Eigenvalue decomposition of TCRen, MJ and Keskin matrices

To assess intrinsic regularity of TCRen potential, we performed eigenvalue decomposition of the corresponding matrix. As this method is applicable only to symmetric matrices, we performed eigenvalue decomposition of its symmetrized form (TCRen.s.cf in **Supplementary Figure 11**).

Eigenvalue decomposition allows to represent all entries of the matrix as:

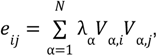

where *λ_α_* are eigenvalues and *V_α_*_,*i*_ are elements of corresponding eigenvectors.

If some eigenvalues prevail and have higher absolute values compared to other eigenvalues, corresponding eigenvectors may be considered as reflecting some properties of residues that govern interaction (in case of TCRen — TCR-peptide recognition).

Eigenvalue spectrum of symmetrized TCRen matrix and correlation coefficients of eigenvectors and AFs are shown in **Supplementary Figure 15, left panel**. For comparison, eigenvalue decomposition was performed also for MJ and Keskin matrices (before analysis they were modified by subtracting mean values <*e_ij_*>) (**Supplementary Figure 15, central and right panels**). In the case of MJ as little as two eigenvalues prevail, both with a strong negative correlation with polarity, that points to the hydrophobic nature of driving forces of protein folding^29^. The form of the eigenvalue spectrum of Keskin matrix is similar to that of MJ matrix, with two dominating eigenvectors negatively correlated with polarity. The notable difference between Keskin and MJ eigenvalue spectrums is that Keskin prevailing eigenvectors also correlate with AF2 (related to secondary structure preferences) and absolute values of other eigenvectors of Keskin are a bit higher compared to MJ matrix, that explains higher correlation of Keskin entries with AF2 of contacting amino acids (**Supplementary Figure 4F**) compared to MJ. However, both entries and prevailed eigenvectors of Keskin have the highest correlation with hydrophobicity AF1, so the major driving force of general protein-protein interactions is also hydrophobic. Eigenvalue spectrum of symmetrized TCRen (**Supplementary Figure 15**) is much more complex than that of MJ and Keskin and a higher number of eigenvalues prevail. Top eigenvectors correlate with different properties of contacting residues: AF4 for eigenvector 1, AF1 for eigenvectors 2 and 3, AF5 for eigenvectors 18 and 19. This analysis suggests that different properties of contacting residues contribute to specificity of TCR-peptide recognition. Although hydrophobicity is important for this process (that is evident from correlation of TCRen entries with products of AF1s of contacting residues in **Supplementary Figure 4F** and correlation of TCRen eigenvectors 2 and 3 with AF1 in **Supplementary Figure 15**) it is not as dominant as in protein folding and protein-protein interaction and other properties (such as electric charge) are also important for TCR-peptide interaction.

Eigenvalue decomposition of asymmetric TCRen matrix can be performed after a simple transformation: residues of the same type in TCR and peptide were considered as distinct entities that resulted in a symmetric matrix of size 38×38 (cysteines are excluded as they are not found in CDR3 regions) with zero values in cells for which both dimensions correspond to residues from the same molecule (TCR or peptide). Eigenvalue spectrum of this 38×38 matrix is shown in **Supplementary Figure 16A**. To examine the relation between eigenvectors and properties of amino acids, correlation coefficients of each eigenvector with 5 AFs were computed separately for residues in TCR and peptide (**Supplementary Figure 16B-C**). Top eigenvectors correlate with different properties of contacting residues and this pattern slightly differs for residues in TCR and peptide side (AF1, AF3, AF4 and AF5 for TCR residues; AF1, AF2 and AF4 for peptide residues; see **Supplementary Figure 16B-C**).

